# “Induction of pancreatic tumor-selective ferroptosis through modulation of cystine import”

**DOI:** 10.1101/827972

**Authors:** Michael A. Badgley, Daniel Kremer, H. Carlo Maurer, Kathleen E. DelGiorno, Ho-Joon Lee, Vinee Purohit, Irina Sagalovskiy, Alice Ma, Jonathan Kapillian, Christina E.M. Firl, Amanda R. Decker, Steve A. Sastra, Carmine F. Palermo, Leonardo R. Andrade, Peter Sajjakulnukit, Li Zhang, Zachary P. Tolstyka, Tal Hirschhorn, Candice Lamb, Tong Liu, Wei Gu, E. Scott Seeley, Everett Stone, George Georgiou, Uri Manor, Alina Iuga, Geoffrey M. Wahl, Brent R. Stockwell, Costas A. Lyssiotis, Kenneth P. Olive

## Abstract

Pancreatic ductal adenocarcinoma (PDA) is the third-leading cause of cancer mortality in the US and is highly resistant to classical, targeted, and immune therapies. We show that human PDA cells are dependent on the provision of exogenous cystine to avert a catastrophic accumulation of lipid reactive oxygen species (ROS) that, left unchecked, leads to ferroptotic cell death, both *in vitro* and *in vivo*. Using a dual-recombinase genetically engineered model, we found that acute deletion of *Slc7a11* led to tumor-selective ferroptosis, tumor stabilizations/regressions, and extended overall survival. The mechanism of ferroptosis induction in PDA cells required the concerted depletion of both glutathione and coenzyme A, highlighting a novel branch of ferroptosis-relevant metabolism. Finally, we found that cystine depletion *in vivo* using the pre-IND agent cyst(e)inase phenocopied *Slc7a11* deletion, inducing tumor-selective ferroptosis and disease stabilizations/regressions in the well-validated KPC model of PDA.

**One Sentence Summary:** Genetic and pharmacological targeting of cystine import induces pancreatic cancer-selective ferroptosis *in vivo*.

## Main Body

The cancer–selective induction of apoptosis through cytotoxic chemotherapy is a foundation of modern oncology. However, some cancers, such as PDA, have proven highly resistant to traditional therapeutic strategies (*1*). In addition to proliferative signals, activating mutations in KRAS (found in >90% of human pancreatic tumors) induce both an increase in the generation of ROS as well as compensatory antioxidant programs (*2–4*). We hypothesize that in this state of elevated ROS flux, ROS detoxification programs become a critical metabolic dependency.

Cellular ROS are neutralized primarily by thiols derived from the semi-essential amino acid cysteine (*5–10*). Antioxidant molecules such as glutathione and thioredoxin serve to position the uniquely– reactive sulfhydryl group of cysteine to catalyze ROS detoxification reactions. As cysteine is ultimately derived from extracellular sources, we examined the effects of culturing PDA cell lines in the absence of cystine, the dimeric, oxidized form of cysteine present in the extracellular environment. In four of five PDA cell lines, cystine withdrawal induced near–complete cell death within 24 hours and this was largely rescued by the lipophilic antioxidant Trolox (**Figure 1A**). Dying, cystine-deprived cells underwent catastrophic destabilization of their plasma membranes, with no visual evidence of nuclear fragmentation (**Supplementary Video 1**).

**Figure 1.**
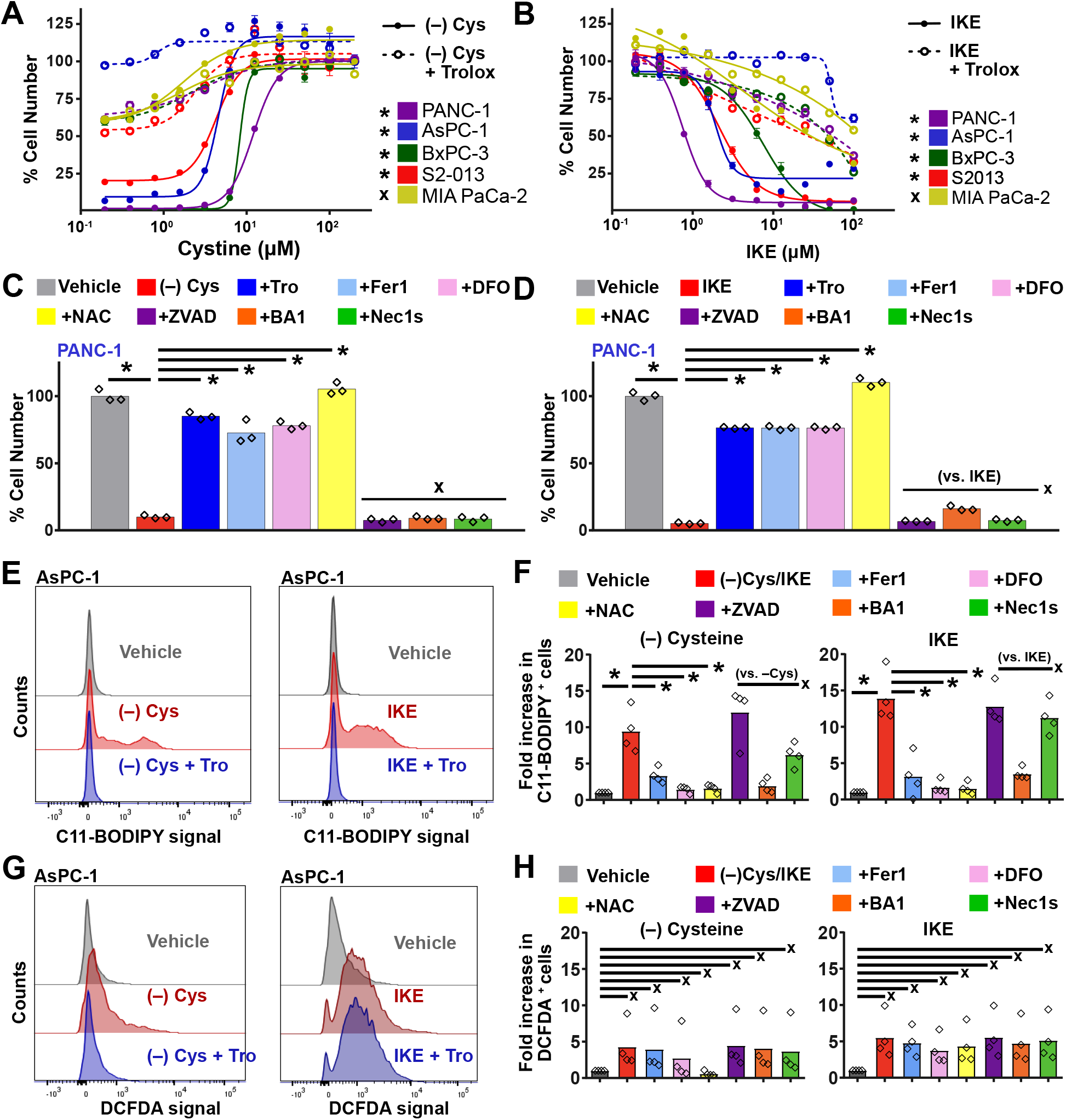
Pancreatic cancer cells require exogenous cystine to avert ferroptosis. **A.** Cystine dose de-escalation curves in PDA cell lines. Cells were cultured in 0 – 200μM cystine in the absence (solid lines) or presence (hashed lines) of Trolox for 24 hours. N=3 biological replicates. Error = +/- SEM. * p< 0.05 comparing maximal cytotoxicity with and without Trolox, Student’s t-test. X = no significant difference. **B.** Dose response curves in PDA cell lines ranging treated with 0 – 100μM IKE in absence (solid lines) or presence (hashed lines) of Trolox for 24 hours. N=3 biological replicates. Error = +/- SEM. * p< .05 comparing maximal cytotoxicity with and without Trolox, Student’s t-test. X = no significant difference. **C.** Viability of Panc1 cells cultured for 24 hours in replete or cystine free media combined with 100μM Trolox (Tro), 500nM ferrostatin-1 (Fer1), 100μM deferoxamine (DFO), 1mM N-acetyl cysteine (NAC), 50μM ZVAD-FMK, 1nM Bafilomycin A1 (BA1) or 10μM Necrostatin 1s (Nec1s). Error bars show +/- SEM. N = 3 biological replicates. * p< .05 by a two-way ANOVA followed by a post-hoc Tukey test, compared to vehicle. * p< .05 by a two-way ANOVA followed by a post-hoc Tukey test, compared to (–) cys. X = no statistically significant difference compared to (–) cys. **D.** Viability of Panc1 cells treated for 24 hours with vehicle or 5μM IKE, alone or in combination with the compounds from (C). Error bars show +/- SEM. N = 3 biological replicates. * p< .05 by a two-way ANOVA followed by a post-hoc Tukey test, compared to vehicle. * p < .05 by a two-way ANOVA followed by a post-hoc Tukey test. X = no statistically significant difference compared to IKE. **E.** Flow cytometry analysis of AsPC-1 cells stained with C11-BODIPY, a marker of lipid oxidative stress, after 6-8 hours of treatment with indicated compounds. Cells were treated with vehicle, no cystine, or no cystine plus 100μM Trolox. **F.** Flow cytometry analysis of PDA cells stained with C11-BODIPY when treated with indicated conditions (cysteine depletion carried out for 8 hours, IKE treatment carried out for 6 hours). Data represented as fold change in number of positive cells. * = p < .05, two-way ANOVA with a post-hoc Tukey test, compared to vehicle. X = no significant difference. **G.** Flow cytometry analysis of AsPC-1 cells stained with H2-DCFDA, a marker of general oxidative stress, after 6-8 hours of treatment with vehicle or no cystine (left panel) or 5μM IKE (right panel), alone or in combination with 100μM Trolox. **H.** Quantification of flow cytometry analysis of PDA cells stained with general oxidation sensor when treated with indicated conditions (cysteine depletion carried out for 8 hours, IKE treatment carried out for 6 hours). Data represented as fold change in number of positive cells. X = no significant difference by one-way ANOVA.

Exogenous cystine is primarily imported by the cystine-glutamate antiporter, system x_c_^−^ (*11*), a heterodimer composed of a common subunit, SLC3A2 (4F2hc), and a specific subunit, SLC7A11 (xCT) (*12*). *We* therefore assessed the effects of pharmacological inhibition of system x_c_^−^ using imidazole ketone erastin (IKE), a potent analog of the system x_c_^−^ inhibitor, erastin (*13, 14*). The effects of system x_c_^−^inhibition closely paralleled that of cystine withdrawal, with near-complete induction of cell death in the same four lines within 24 hours **(Figure 1B)**. Cell death induced by system x_c_^−^inhibition was rescued by Trolox (**Figure 1B**) and appeared identical to that of cystine starvation, but distinct from staurosporine–induced apoptosis (**Supplementary Figure 1A; Supplementary Videos 2, 3**). Neither cystine withdrawal nor system x_c_^−^ inhibition (collectively referred to as “cysteine depletion”) led to caspase 3 cleavage (**Supplementary Figure 1B**), indicating a form of non-apoptotic cell death.

The rapid, oxidative cell death resulting from cysteine depletion was reminiscent of ferroptosis, a form of iron–dependent, non-apoptotic cell death previously associated with erastin treatment (*15*). Indeed, near-complete rescue of cysteine-depletion induced cell death was observed following co-treatment with either deferoxamine (DFO, an iron chelator) ferrostatin-1 (Fer1, a ferroptosis inhibitor) or N-acetyl cysteine (NAC, a cell permeable analog of cysteine). By contrast, the apoptosis inhibitor ZVAD-FMK had no effect, and inhibitors of necroptosis and autophagy offered only moderate rescue in a subset of lines, consistent with previous reports (*16, 17*) (**Figure 1C, D, Supplementary Figure 1C–E**).

Another defining feature of ferroptosis is the rapid accumulation of lipid ROS. Following cysteine depletion, we observed a sharp increase in oxidation of the sensor dye C11-BODIPY preceding the onset of cell death by at least an hour (**Figure 1E, Supplementary Figure 2**) and this was fully rescued by cotreatment with Trolox and other agents that rescued cysteine depletion-induced cell death (**Figure 1E,F**). By contrast, elevated general ROS from cysteine depletion was not rescued by Trolox, arguing against a more generalized oxidative response **(Figure 1G, H)**. We conclude that cysteine depletion leads to the induction of ferroptosis in the majority of human PDA lines tested.

To assess the relevance of system x_c_^−^ to human patients, we examined the expression of the specific subunit *SLC7A11* across six public PDA datasets that included matched normal tissues. A meta-analysis found a roughly 1.5–fold increase in *SLC7A11* RNA expression in PDA vs. normal pancreas tissue (p= 4.1 x 10^−7^, random effects model) (**Supplementary Figure 3A**). Functionally, *SLC7A11* expression in these cohorts was significantly associated with signatures of redox stress (**Supplementary Figure 3B-D**). Based on the predominant expression of *SLC7A11* in the epithelium of PDA compared to stroma in laser capture microdissected human PDA samples (**Supplementary Figure 4A**) (*18*), these functions likely reflect its role in the malignant compartment.

Next, we examined expression data from The Cancer Genome Atlas (TCGA) and found that pancreatic tumors exhibit high absolute expression for *SLC7A11* compared to other cancers, with comparatively low variance (**Supplementary Figure 4B**). We also found that *SLC7A11* expression was significantly upregulated relative to normal tissues in many tumor types (**Supplementary Figure 4C**). Among those cancers with upregulation of *SLC7A11*, its expression was frequently associated with reduced overall survival (**Supplementary Figure 4D**). *SLC7A11* expression is therefore associated with malignancy, oxidative processes, and poor outcome across a wide range of human cancers.

To date, *in vivo* evidence for the role of system x_c_^−^ in cancer has largely relied on *ex vivo* assays, direct injection of chemical probes (*19*), or implantation of cells with existing alterations that could impact engraftment (*20*). In order to determine the reliance of established pancreatic tumors on system x_c_^−^ in a physiological context, we employed a dual recombinase genetic engineering strategy based on the well–validated “KPC” mouse (*21*). *<u>K</u>ras^FSF.G12D/+^; <u>p</u>53^R172H/+^; Pdx1<u>F</u>lpO^tg/+^; <u>S</u>lc7a11^Fl/Fl^; <u>R</u>osa26^CreERT2/+^* (KPFSR) mice (see **Supplementary Figure 5** for more information) developed tumors histopathologically identical to those from KPC mice: typically a single, focal mass with moderate differentiation and robust stromal desmoplasia (**Supplementary Figure 6A**), in the context of surrounding premalignant histopathology (**Supplementary Figure 6B**). Following the identification of 4–7mm diameter pancreatic tumors by ultrasound, KPFSR mice were randomized to receive 6 daily doses of either vehicle or tamoxifen in order to induce genetic recombination to systemically delete both copies of *Slc7a11*. PCR performed on normal and tumor tissues confirmed at least partial recombination *in vivo* following tamoxifen delivery (**Supplementary Figure 6C**). We found that acute deletion of *Slc7a11* in tumor-bearing KPFSR mice nearly doubled median survival (**Figure 2A**, p< 0.0295, Log Rank) and a post-hoc analysis confirmed similar tumor sizes at enrollment (**Supplementary Figure 6D**). Most tamoxifen–treated tumors exhibited a period of stable disease or regression whereas no such responses were observed in vehicle-treated animals (**Figure 2B, Supplementary Figure 6E**). Critically, both the extension of overall survival and induction of tumor responses were completely reversed through the addition of NAC to the drinking water of tamoxifen-treated animals (**Figures 2A,B**), confirming these phenotypes were related to cysteine metabolism. Modeling of tumor growth rates (*22*) showed that tamoxifen-treated tumors grew significantly slower than those in vehicle-treated mice (**Supplementary Figure 6F**). Notably, one tumor regressed until it was undetectable by ultrasound (**Supplementary Figure 6G**); this animal later succumbed to a new tumor arising in a different site in the pancreas. While some recombination was still evident in tumors at necropsy, Western blotting detected similar total levels of Slc7a11 protein in tamoxifen- and vehicle-treated tumors at endpoint, suggesting the outgrowth of unrecombined tumor cells (**Supplementary Figure 7A,B**).

**Figure 2.**
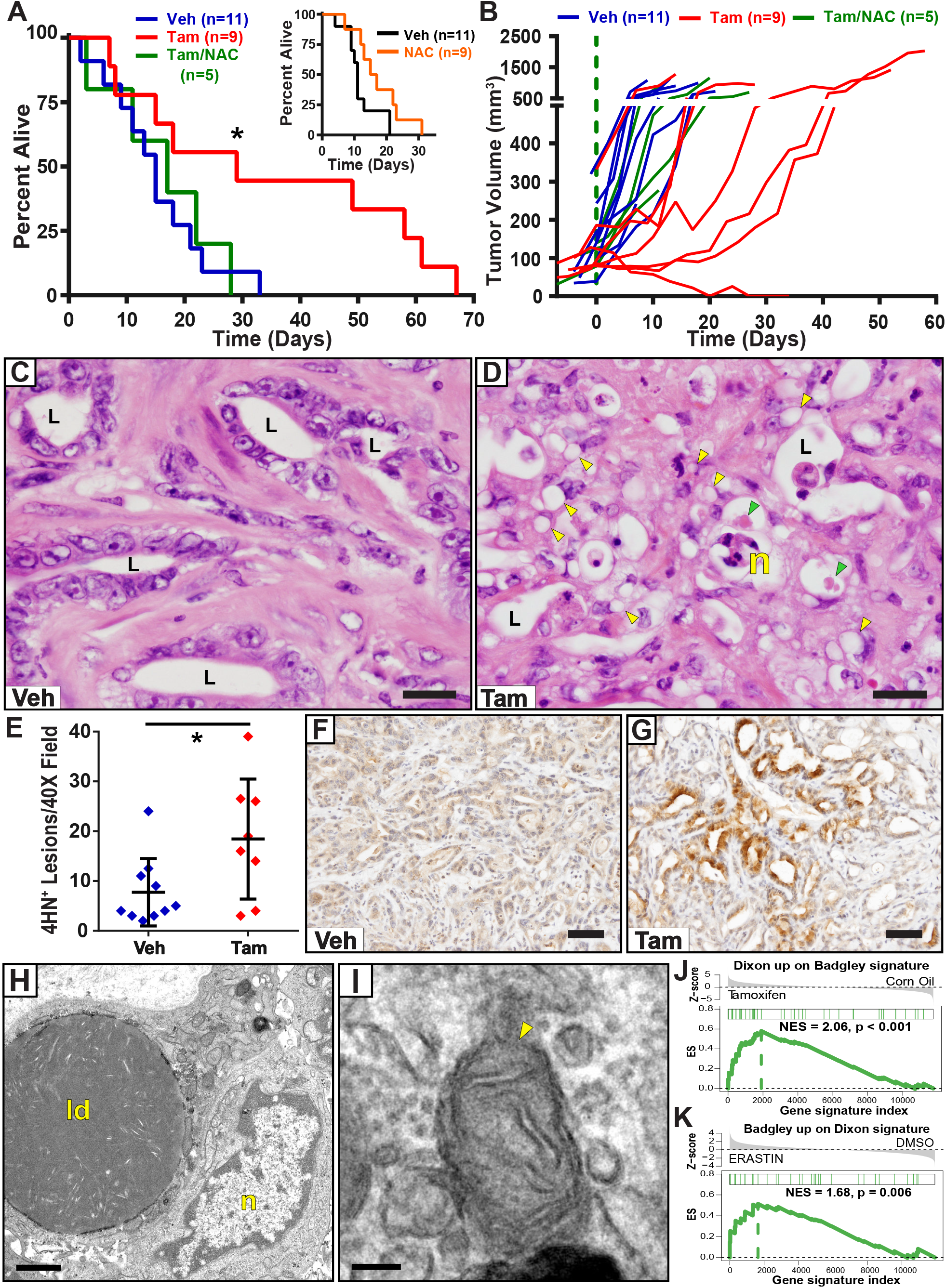
Deletion of Slc7a11 extends survival in a mouse model of pancreatic cancer through ferroptosis. **A.** Kaplan-Meier survival curve of KPFSR mice treated with Vehicle (N = 11, median survival 15 days), Tamoxifen (N = 9, median 29 days), or Tamoxifen/NAC (N = 5, median 17 days). * p< .0295, Log-rank. INSET: Survival curves of KPC mice treated with NAC alone versus historical controls are shown for comparison. **B.** Growth curves derived from 3D ultrasound data for each tumor following enrollment on Day 0. **C, D.** Histology images of KPFSR pancreatic tumors treated with vehicle (C) or tamoxifen (D) to induce deletion of *Slc7a11*. treated versus untreated tumors. Vehicle treated tumors exhibit a typical appearance for this model. L = lumen of malignant epithelium. Tamoxifen–treated tumors exhibit a distinctive histolopathology with subcellular inclusions (yellow arrows), megamitochondria (green arrows), and cellular necrosis (n). Scale = 20μm. **E.** Quantification of the number of 4-hydroxynonenal (4HN) lesions in per low powered image. * p< .05, Student’s t-test. **F, G.** Representative images of IHC for 4-hydroxynonenal (4HN). Scale = 50μm. **H.** Transmission electron microscopy images from a KPFSR tumor treated with tamoxifen to induce *Slc7a11* deletion. ld = lipid droplets, n= nucleus. Scale = 1μm. **I.** TEM images of aberrant mitochondria in *Slc7a11* deleted tumor tissue. Yellow arrows indicate disrupted mitochondrial membranes and cristae. Scale = 100nm. **J.** Gene set enrichment analysis showing enrichment of a previously published ferroptosis expression signature from erastin-treated HT1080 cells (Dixon) compared to genes that are differentially expressed between *Slc7a11* deleted vs. unrecombined PDA cells (Badgley). The Badgley signature was generated by laser capture microdissection and RNA-sequencing of tamoxifen and vehicle-treated KPFSR tumors. **K.** Reciprocal comparison from (J), showing enrichment of the Badgley signature in the Dixon gene set.

Prior reports have relied on marker genes or metabolites extracted from tumor tissues to argue for the induction of *in vivo* ferroptosis (*13, 19*). The absence of a histopathological characterization of tumor ferroptosis has hindered our ability to link individual markers to non-apoptotic cell death in a physiological context. A histological examination of KPFSR mice found no aberrant pathology in the non–pancreatic tissues of tamoxifen–treated animals. However, within their pancreatic tumors, we observed lesions with an unusual cellular phenotype characterized by ballooned epithelial cells with cytoplasmic vacuolization and occasional megamitochondria (**Figure 2C, D; Supplementary Figure 7C)**. These lesions preferentially occurred in juxtaposition to areas of frank necrosis, yet they were negative for the apoptosis marker cleaved caspase 3 and exhibited no noticeable change in proliferation markers **(Supplementary Figure 7D, E)**. Careful examination of vehicle-treated KPFSR tumors as well as tumors from the classic “KPC” model of PDA identified occasional, limited examples of this phenotype, typically in association with frank necrosis.

Notably, we observed focal accumulation of 4-hydroxynonenal (4HN), a byproduct of lipid peroxidation, by IHC staining in tamoxifen-treated KPFSR tumors (**Figure 2E**), particularly in association with the vacuolized lesions (**Figure 2F,G**). Transmission electron microscopy (TEM) on a tamoxifen-treated KPFSR tumor identified the vacuolized structures as abnormally large lipid droplets (**Figure 2H)** and this was confirmed by Oil Red O staining (**Supplementary Figure 7F)**. TEM also revealed frequent examples of aberrant mitochondria in the malignant epithelial cells, with disrupted cristae and loss of mitochondrial membrane integrity (**Figure 2I**), matching prior *in vitro* reports on the effects of ferroptosis on mitochondria (*15, 23, 24*).

Finally, to directly compare the response of the *Slc7a11*–deleted tumors to the *in vitro* studies that have classically been used to define the phenomenon of ferroptosis, we performed laser capture microdissection and RNA-sequencing on malignant epithelial cells from KPFSR tumors treated with vehicle or tamoxifen. We found that genes that are differentially expressed in *Slc7a11*–deleted PDA cells *in vivo* are enriched for the ferroptosis signature defined from erastin-treated HT-1080 cells in vitro (*25*), and vice versa (**Figure 2J,K**). By contrast, none of the ten Gene Ontology (GO) apoptosis gene sets were enriched in tamoxifen-treated KPFSR tumors (**Supplementary Table 1**). Taken together with our earlier findings, we conclude that the novel phenotype observed in tamoxifen-treated KPFSR tumors is a histologically identifiable *in vivo* manifestation of ferroptosis.

We next examined the mechanisms of ferroptosis in PDA cells (*13, 26–28*). Prior studies have indicated that cysteine-depletion–induced ferroptosis occurs due to the loss of glutathione, which serves as a cofactor for the critical lipid peroxide detoxifying enzyme GPX4 (*13, 29*). Consistent with this model, we observed a significant loss of intracellular cysteine and a near-complete loss of glutathione after 6 hours of IKE treatment in two human PDA lines (**Figure 3A, Supplementary Figure 8A**). Moreover, provision of an alternative glutathione source via the cell permeable analog glutathione ethyl ester (GSH-EE) led to complete rescue of cysteine-depletion induced ferroptosis (**Figure 3B, Supplementary Figure 8B,C**). This was associated with complete reversal of lipid peroxidation (**Figure 3C**) but only partial reversal of general ROS induction (**Figure 3D**), continuing the close association of lipid peroxide induction with ferroptosis. However, inhibition of glutathione biosynthesis using buthionine sulfoximine (BSO, **Supplementary Figure 8D**) did not induce lipid ROS (**Figure 3E**) and had no impact on cell viability (**Figure 3F**). This result highlights contradictory findings regarding the sufficiency of glutathione depletion for the induction of ferroptosis in different cell lines (*13, 30*). We inferred that in PDA cells, the loss of one or more additional cysteine–derived metabolites may be critical to the induction of ferroptosis.

**Figure 3.**
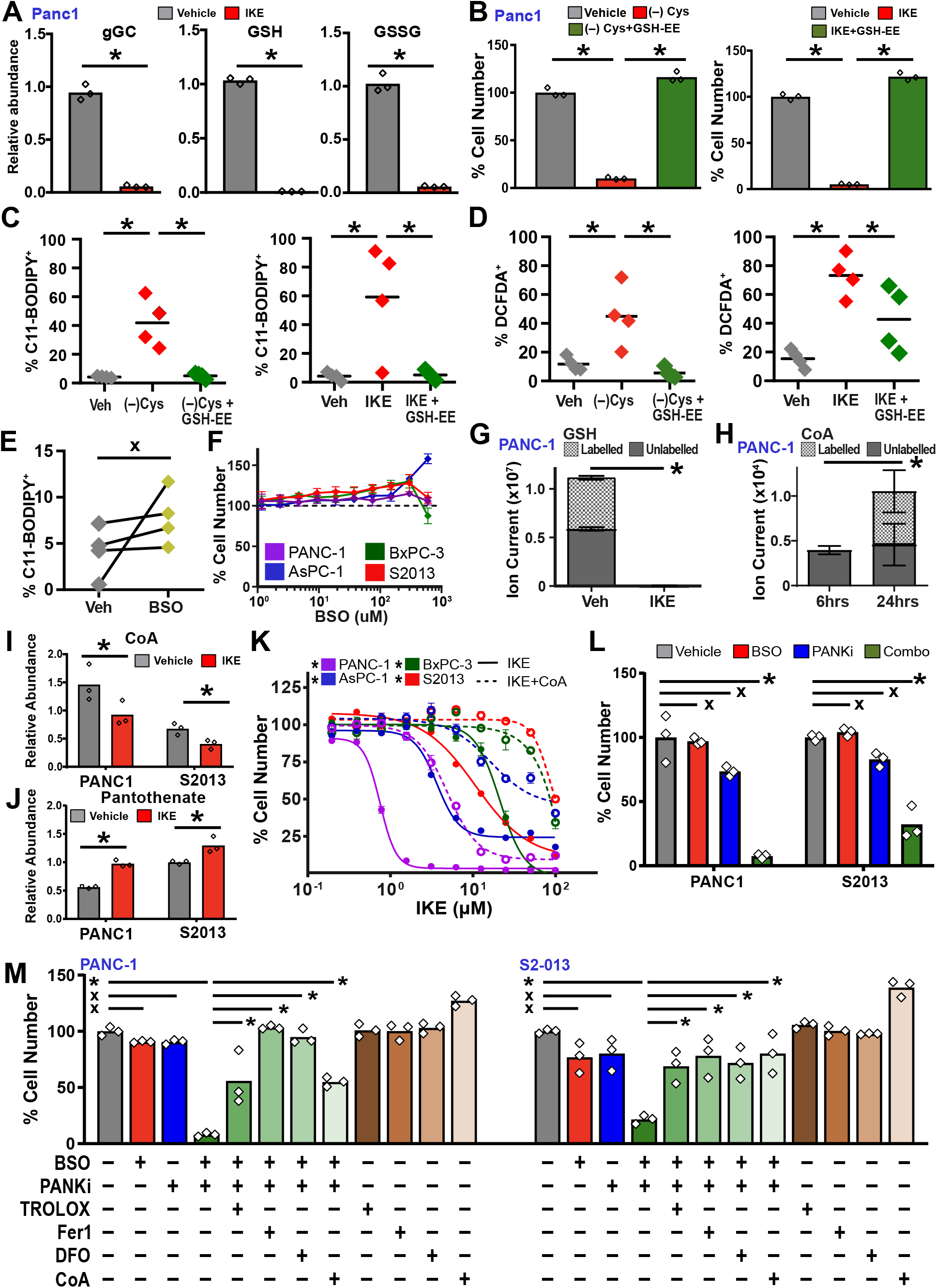
GSH depletion is not sufficient to induce ferroptosis. **A.** The effects of system x_c_^−^ inhibition on the depletion of intracellular cysteine (gGC), GSH, and oxidized GSH (GSSG) in PANC-1 and S2-013 cells, as measured by (LC-QqQ-MS). Analytes were extracted and processed 6 hours after treatment initiation. Cells were treated with vehicle (0.05% DMSO) or 5 μM IKE. N = 3 biological replicates, error bars +/- SD. * p< .05 by Student’s t-test. X = not significant. **B.** Viability of PDA cells grown in the absence of cystine or when treated with the system x_c_^−^ inhibitor IKE by supplementing with GSH-ethyl esther (GSH-EE). N = 3 biological replicates, error bars = +/- SD. * p< .05 by Student’s t-test. Measurements are taken 24 hours after treatment initiation. X = not significant. **C, D.** Flow cytometry analysis of PDA cells stained with C11-BODIPY (C) or H2DCFDA (DCFDA) after 6 hours of treatment with vehicle (0.2% DMSO), IKE (10 μM), no cystine, and/or GSH-EE (1mM). * p < 0.05 by one-way ANOVA with post hoc Tukey test. X = not significant. **E.** LC-QqQ-MS measurements of intracellular GSH in PANC1 cells treated with vehicle (0.2% DMSO) or 150μM BSO for 6 hours or 24 hours. N = 3 biological replicates, error bars = +/- SD. * p < .05 by Student’s t-test. **F.** Dose response curves in PDA cells treated for 24 hours with BSO. N = 3 technical replicates, error bars = +/- SD. **G.** LC-TOF-MS analysis of GSH in ^13^C-cystine labelled PANC-1 cells treated with vehicle (.1% DMSO) or IKE (5 μM) for 6 hours. **H.** The incorporation of ^13^C labeled cystine into intracellular CoA pools as measured by LC-TOF-MS. Cells were cultured for 6 or 24 hours (as indicated on x-axis) prior to sample collection. Labeled metabolite signal indicated by gridded pattern. N = 3 biological replicates, error bars = +/- SD. **I.** LC-QqQ-MS measurements of CoA levels in PDA cells treated with vehicle and IKE for six hours. **J.** LC-QqQ-MS measurements of pantothenate levels in PDA cells treated with vehicle and IKE for six hours. **K.** Dose response curves of sensitive cell lines treated with IKE (solid lines) and in combination with CoA (200 μM, dotted lines). * p < 0.05 comparing maximal cytotoxicity in CoA treated vs. untreated conditions for each line, Student’s t-test. **L.** Viability of PDA cells after treatment with vehicle, BSO, PANKi, and combo for 24 hours. * p<0.05, oneway ANOVA with Tukey’s. **M.** Viability of PDA cells treated with the indicated combinations of BSP, PANKi, Trolox, Fer1, DFO, or CoA.

To test this, we treated human PDA cells with ^13^C-labelled cystine and examined the synthesis of cysteine-derived metabolites from exogenous cystine pools, before and after treatment with IKE. As expected, exogenous labelled cysteine was rapidly incorporated into glutathione pools and this flux was immediately inhibited by treatment with IKE (**Figure 3G**). By contrast, we did not detect significant contribution of exogenous cystine into taurine, lactate, citrate, or glutamate over the course of 24 hours (**Supplementary Figure 8E**). We did, however, observe significant labelling of coenzyme A (CoA) pools (**Figure 3H**), an important cofactor that is synthesized from cysteine via the pantothenate pathway. CoA is a precursor to mevalonate, cholesterol, and coenzyme Q_10_ (CoQ_10_), among other metabolites, and is required for fatty acid synthesis, fatty acid β-oxidation, and many other critical metabolic processes. Consistent with flux of cysteine into CoA, we found that inhibition of system x_C_^−^ reduced total CoA levels (**Figure 3I**) and simultaneously increased levels of pantothenate (**Figure 3J**), a metabolite upstream of cysteine incorporation in the CoA synthesis pathway.

To directly assess whether CoA plays a role in cysteine–depletion–induced ferroptosis, we treated PDA cells with exogenous CoA in the context of system x_C_^−^ inhibition; while CoA itself cannot cross the plasma membrane, it is degraded extracellularly to the membrane–permeable precursor 4’-phosphopantetheine and then restored to CoA within cells by Coenzyme-A Synthase (*31*). Strikingly, exogenous CoA rescued all four PDA lines from IKE-induced ferroptosis (**Figure 3K**), implying a critical role for CoA metabolism in the regulation of ferroptosis. We next examined whether depletion of CoA could impact ferroptosis in PDA cells. We utilized PANKi, an inhibitor of pantothenate kinase (PANK) enzymes which catalyze the first step of CoA synthesis from pantothenate (*32*) (**Supplementary Figure 9A**). As expected, treatment of PDA cells with PANKi resulted in increased levels of pantothenate (**Supplementary Figure 9B**). Notably, we found that PANK inhibition sensitized PDA cells to system x_C_^−^ inhibition, showing an interaction between CoA synthesis and cysteine depletion (**Supplementary Figure 9C,D**). To assess whether inhibiting both the GSH and CoA arms of cysteine metabolism is sufficient for induction of ferroptosis, we treated PDA lines with the combination of BSO and PANKi. Critically, while neither agent affected PDA cell viability alone, the combination potently synergized to reduce cell viability (**Figure 3M**). Cell death from combination BSO/PANKi treatment was rescued by Fer-1 and DFO confirming induction of ferroptosis (**Figure 3M**). We also found cell death was rescued by idebenone (a membrane permeable analog of CoQ10) and by oleic acid, a monounsaturated fatty acid that lacks bis-allylic sites that can initiate lipid peroxidation (*19*). By contrast, a saturated fatty acid (palmitic) and a polyunsaturated fatty acid (linoleic) both failed to rescue ferroptosis (**Supplementary Figure 9E,F**). Together, these data indicate that multiple cysteine-derived metabolites contribute to the regulation of ferroptosis, including both glutathione and Coenzyme A (**Supplementary Figure 9G**).

Finally, we sought to leverage these findings as a means to selectively target pancreatic tumors for ferroptotic cell death. While potent system x_C_^−^ inhibitors are not yet ready for translation, an engineered enzyme, cyst(e)inase, was recently reported that effectively degrades both cystine and cysteine in mammalian circulation (*33*). Cyst(e)inase is stable in mammals for days and is being developed clinically to treat the metabolic disorder cystinuria. We hypothesized that that cyst(e)inase could therapeutically recapitulate the effects of cystine starvation in PDA. *In vitro*, cyst(e)inase treatment reduced cell viability in all four sensitive PDA lines, albeit with slower kinetics than system x_C_^−^ inhibition (**Figure 4A, Supplementary Figure 10A**). The loss of viability was rescued by ferroptosis inhibitors Fer1 and DFO (**Figure 4B, Supplementary Figure 10B**). Treatment was also associated with induction of lipid oxidation, as measured by C11-BODIPY staining (**Figure 4C, Supplementary Figure 10C**), leading to the conclusion that cyst(e)inase can induce ferroptosis in PDA cell lines.

**Figure 4.**
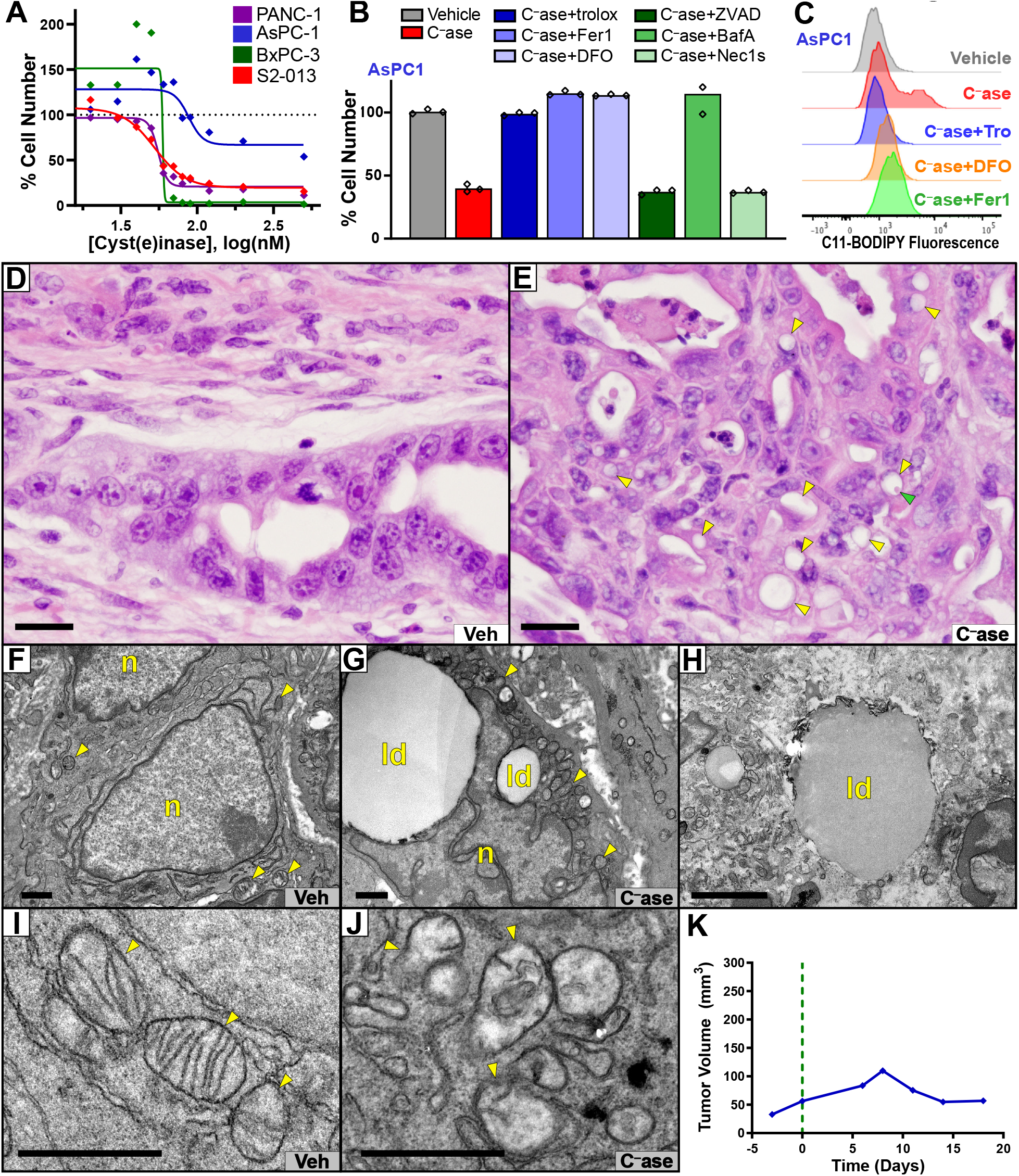
Cyst(e)inase treatment induces tumor-selective ferroptosis in KPC mice. **(A)** Viability dose response curves for PDA cells treated with increasing concentrations of cyst(e)inase for 48 hours (Aspc1) or 72 hours (Panc1, BxPC3, S2-013). N=3 biological replicates. **(B)** Viability of AsPC1 cells treated with cysteinase for 72 hours, alone and in combination with rescue agents (Trolox, Fer1, and DFO) or other cell death inhibitor (ZVAD-FMK, Bafilomycin A1, and Necrostatin-1s). **(C)** Lipid oxidation in AsPC1 cells measured by C11-BODIPY flow cytometry showing induction of lipid oxidation by cyst(e)inase and rescue by inhibitors of ferroptosis. **(D,E)** H&E stained histopathology of pancreatic tumors from the KPC model, treated with vehicle (D) or cyst(e)inase (E). Yellow arrowheads indicate lipid droplets. Green arrowhead indicates megamitochondrion. Bars= 20μm. **(F-J)** Transmission electron microscopy of pancreatic tumors from the KPC model treated with vehicle (F,I) or cyst(e)inase (G,H,J). Yellow arrowheads indicate mitochondria; ld= lipid droplets; n= nucleus. Cyst(e)inase treated tumor cells exhibited frequent, large lipid droplets and large numbers of aberrant mitochondria with ruptured mitochondrial membranes and disrupted cristae. (J) shows a large extracellular lipid droplet from a cyst(e)inase treated tumor. Bars= 1μm. **(K)** Tumor growth curve derived from 3D ultrasound on a KPC mouse treated with cys(e)inase on day 0 (green line) showing a tumor regression in response to treatment. Similar tumor responses have not been previously observed vehicle-treated KPC tumors.

To determine whether cyst(e)inase could induce ferroptosis in pancreatic tumors *in vivo*, we performed a short-term intervention study in KPC mice, a gold-standard genetically engineered model of PDA that is highly resistant to therapeutic intervention (*21, 34*). Tumor-bearing KPC mice identified by ultrasound were treated for 10 days with either a low dose (50mg/kg, q3d, iv) or a high dose (100mg/kg, q2d, iv) of cyst(e)inase, or a vehicle control (n=2 per group). Four additional KPC mice were treated with low-dose cysteinase and monitored longitudinally by 3D high resolution ultrasound. Across all eight cyst(e)inase-treated animals, we observed a striking histpathological recapitulation of the lesions observed in tamoxifen-treated KPFSR mice, though greater in both degree and penetrance (**Figure 4D,E; Supplementary Figure 11A-B**). Numerous cells with large lipid droplets were identified, often in juxtaposition to areas of cell death (**Supplementary Figure 11C**). In some cases, largescale tissue disruption and decompressed blood vessels were noted (**Supplementary Figure 11D**). Necrosis was observed both in focal clusters (**Supplementary Figure 11E**) and in large fields (**Supplementary Figure 11F**), and was particularly apparent in the two tumors treated with high-dose cyst(e)inase (**Supplementary Figure 11G**). TEM performed on necropsy tumor samples from the short-term study showed medium to large lipid droplets in all but one of the cyst(e)inase treated tumors (**Figure 4F,G**), and we observed several examples of extracellular lipid droplets, presumably extruded following cell death (**Figure 4H**). Moreover, TEM analysis identified widespread mitochondrial structural defects in all cyst(e)inase treated tumors, but not in vehicle-treated tumors (**Figure 4I,J**). Immunohistochemistry performed on necropsy tumor tissues showed focal accumulation of 4HN in the absence of cleaved caspase 3 positivity in lipid droplet–rich lesions (**Supplementary Figure 12A**). Finally, reconstruction of longitudinal 3D ultrasound data revealed tumor stabilizations or regressions in all four KPC mice treated with cyst(e)inase, whereas historical vehicle-treated controls never stopped growing (**Figure 4K, Supplementary Figure 12B**). Normal tissues that were unaffected by pancreatic cancer in KPC mice showed no additional histopathology from cyst(e)inase–treatment. Thus we conclude that the therapeutic depletion of cysteine/cystine can lead to the selective induction of ferroptosis in Kras/p53 mutant autochthonous pancreatic ductal adenocarcinomas.

Since it was first described in 2012 (*15*), ferroptosis has been investigated as a potential contributor to human pathology, notably neurodegenerative diseases, traumatic brain injury, acute kidney injury, and cancer (*35*). However, *in vivo* evidence for ferroptosis has been limited as current ferroptosis–inducing agents have poor pharmacological properties and there are no established, specific markers of ferroptosis *in vivo*. Using both genetic and pharmacological models, we observed a novel histopathology in response to cysteine depletion in established Kras/p53 mutant pancreatic tumors, which we identified as an *in vivo* manifestation of ferroptosis.

All cells, whether normal or malignant, must carefully manage the disposition of atoms, energy, and electrons. Cancer cells selectively prioritize the production of biomass in support of persistent cell division, generally through pathways that incur an electron imbalance and thereby generate ROS (*36*). Cysteine-derived metabolites such as glutathione play a key role in managing electron imbalances that could otherwise lead to ferroptosis. However, GSH depletion alone is not sufficient to cause ferroptosis in PDA cells, implicating additional cysteine-derived metabolites in the management of ferroptosis. The identification of coenzyme A as a key downstream cysteine-derived metabolite that contributes to the control of ferroptosis provides a direct link to pathways that control the production, breakdown, and repair of lipid membrane components. While additional work will be necessary to illuminate precisely how CoA– dependent processes contribute to suppression of ferroptosis, one potential model is that inhibition of GSH elevates the amount of cysteine available for the production of CoA, and by extension downstream metabolites that regulate detoxification of lipid ROS species. This model is strongly supported by the previous discovery of the ferroptosis inducer FIN56 (*37*), which inhibits synthesis of mevalonate-derive products such as CoQ10, and the recent identification of a glutathione-independent suppressor of ferroptosis, FSP1, which catalyzes the reduction CoQ10 to directly quench lipid peroxyl radicals (*38, 39*).

Apoptosis is the predominant means by which many clinical anticancer agents are known to kill tumor cells. However, traditional chemotherapy has had limited efficacy in PDA and other forms of cell death have not yet been leveraged as effectively in the clinic. The cancer–selective induction of ferroptosis following systemic deletion of *Slc7a11* or cyst(e)inase treatment demonstrates the potential therapeutic utility of targeting cysteine metabolism. Cysteine depletion was remarkably well tolerated in non-tumor tissues. We speculate that this is due to the elevated ROS flux in KRAS-driven PDA cells, combined with the highly hypoxic microenvironment that arises in these poorly perfused–tumors (*34*). The general tolerability of acute *Slc7a11* deletion is consistent with prior reports that germline *Slc7a11* knockout mice are viable and healthy into adulthood, though particularly sensitive to redox stress (*40*), and cyst(e)inase was previously shown to be well tolerated in healthy mammals (*33*). Combined with evidence that this pathway is upregulated across human cancers, we consider cysteine metabolism to be a strong target for clinical development, both for PDA and possibly other malignancies. Taken together, our results underscore an emerging understanding that maintenance of intracellular redox state is critical to cancer cell viability and may be targeted as form of metabolic anticancer therapy.

## Materials and Methods

### Cell lines, cell culture, and viability assays

All cell lines were obtained from ATCC and tested negatively for mycoplasma infection. Cells were maintained under standard conditions at 37°C and 5% CO2. Cells were grown in DMEM (Life Technologies, 12430-054) supplemented with penicillin and streptomycin (Corning, 30-003-CI), 10% FBS (Life Technologies, 10438-034), and MEM NEAA (Life Technologies, 11140-050), unless otherwise indicated. For cystine withdrawal experiments, cells were cultured in DMEM lacking glutamine, methionine, and cystine (Life Technologies, 21013-024), supplemented with 200mM methionine (Sigma, M9625), 4mM glutamine (Life Technologies, 25030-081), 10% FBS, penicillin and streptomycin and varying concentrations of cystine (Sigma, C8755), depending on the given experiment. For all cell viability experiments, cells were plated in a 96-well plate format. PANC-1 and S2-013 cells were plated at 4,000 cells per well; MIA PaCa-2, BxPC-3, and AsPC-1 cells were plated at 8000 cells per well. Cells were allowed to seed overnight, and subsequently treated with compounds at indicated concentrations and for indicated lengths of time. The following is a list of all chemical compounds used in cell culture experiments: Trolox (Sigma, 238813), ZVAD-FMK (SelleckChem, S7023), Bafilomycin A1 (Sigma, B1793), Necrostatin-1s (BioVision, 2263-1), Ferrostatin-1 (Sigma, SML0583), N-acetyl-L-cysteine (Sigma, A9615), buthionine sulfoxamine (Sigma, B2515), deferoxamine (Sigma, D9533), EIPA (Sigma, A3085), propargylglycine (Sigma, 81838), GSH-ethyl ester (Sigma, G1404), staurosporine (Sigma, S4400), PE and IKE (provided by Brent Stockwell), CoA (Sigma, C3144). Cyst(e)inase was provided by Everett Stone and George Georgiou. For PANKi and BSO combination studies, the following compounds were used: Pantothenate kinase inhibitor (Calbiochem, 537983), L-Buthionine-(S,R)-Sulfoximine (Cayman,14484), Imidazole ketone erastin (Medchem express, HY-114481), Ferrostatin-1 (Cayman, 17729), Trolox (Cayman, 10011659), Deferoxamine (mesylate) (Cayman, 14595), Coenzyme A (Cayman, 16147), Glutathione ethyl ester (Cayman, 14953), Palmitate (calcium salt) (Cayman, 10010279), Oleic Acid (Cayman, 90260), Linoleic Acid (Cayman, 90150), DMSO (Sigma, D2650), Ethanol (Decon labs, 2701), and Chloroform sterile filtered (Sigma, C2432).

Compounds were formulated according to the manufacturer’s instructions. For viability assays testing PANK inhibition and combinatorial BSO treatment, cells were plated in a 96-well plate format at 1,000 cells per well, allowed to seed overnight, then treated with compounds at indicated concentrations and for indicated lengths of time. These viability assays utilized the Cell-Titer-Glo 2.0 reagent (Promega, G9243) according to the manufacturer’s instructions. Briefly, all wells from 96-well plates were aspirated followed by the addition of 100 μL of Cell-Titer-Glo 2.0 reagent to each experimental well. Plates were gently agitated for 10 minutes to promote adequate mixing. Luminescence was subsequently measured using a SpectraMax M3 plate reader. All other viability assays were carried out using the AlamarBlue reagent (Thermofisher, 88952) according to the manufacturer’s instructions. In brief, 10 μL of AlamarBlue reagent was added each well containing 100 μL of experimental media. Plates were gently agitated for 1 minute to promote adequate mixing. Once wells had reached uniform color, plates were incubated at standard culture conditions indicated above for 1-4 hours. Plates were subsequently assessed for fluorescent readout on a Promega plate reader using the green filter.

### Light and fluorescent microscopy

Still transmitted light images were captured on an Olympus CKX41. Time lapse images and fluorescent microscopy still images were captured on a Nikon A1RMP. For time lapse videos, one image was captured every minute for up to 24 hours post treatment initiation. Cells were plated in a 6-well plate format, (PANC-1: 250,000 cells per well). For fluorescent ROS detection, cells were stained with C-11 BODIPY (Invitrogen, C10445) at a concentration of 2μM for 30 minutes prior to time lapse imaging. Cells were washed with PBS three times and incubated in Live-Cell Imaging Solution (Life Technologies, A14291DJ) during the duration of live-cell imaging. To detect lipid ROS at static time points by fluorescence microscopy, cells were plated in 1u-Slide 8 well ibiTreat dishes (ibidi, 80827). PANC-1 cells were plated at 30,000 cells per chamber. Cells were seeded overnight and subsequently subjected to indicated treatments for indicated times. Upon experiment completion, cells were visualized on a Nikon A1RMP in the red, green, and transmitted light channels. Images from all three channels were then overlaid to produce final images.

### Flow cytometric detection of ROS

Cell lines were plated in quadruplicate at the cell numbers indicated previously in a 96-well plate format. Cells were allowed to seed overnight and were subjected to various compound treatments for indicated times. Cells were then incubated for 30 minutes in live-cell imaging solution containing the pertinent ROS dye at the following concentrations: C-11 BODIPY, 2 μM; and H2DCFDA, 10 μM (Invitrogen, C6827). Cells were then washed with PBS, trypsinized with .25% trypsin (Life Technologies, 25200-056), and neutralized with 10% FBS in PBS at a 1:1 volume. Cells were strained through a 40μM strainer (BD Falcon, 08-771-1), and analyzed on a MACSQuant Analyzer 10, BD LSRII, or a BD Fortessa with high throughput attachment depending on the application. A minimum of 4,000 cells were analyzed per condition. For both C-11 BODIPY and H2DCFDA, signal was analyzed in the FITC channel. Software analysis was carried out using FlowJo V10.

### Cell-based Thermal Shift Assay

SKBR3 cells were seeded in 3 T-175 flasks, 10 million cells per flask, overnight, then treated with DMSO (vehicle), 20μM PANKi, or 20μM SGC-GAK-1 for two hours, trypsinized, and washed with PBS. Cell pellets were resuspended with 230 μL of PBS + protease inhibitors (Roche) and split into ten 20μL PCR tubes. Cells were heated to indicated temperatures for 3 minutes in two batches in a preheated thermocycler, then placed at room temperature for three minutes, then snap frozen in liquid nitrogen. Samples were placed into the PCR machine only when it reached the desired temp, and heated for three minutes, following by three minutes at RT. Cells were lysed via two freeze-thaw cycles of liquid nitrogen/25°C, vortexed, and then spun at 20,000 rpm for 20 min at 4°C. Supernatant was removed, sample buffer added, and samples boiled heated to 70°C for 10 min, and then loaded for gel electrophoresis at 200V for 45min. Western blotting for PANK1 was performed according to standard protocols.

### Cyst(e)inase *in vitro* studies

Cells were plated in a 96-well plate format as follows: PANC-1 and S2-013 cells were plated at 2,000 cells per well; MIA PaCa-2, BxPC-3, and AsPC-1 cells were plated at 3000 cells per well. Cells were allowed to grow overnight, and subsequently treated with 90nM cyst(e)inase in the presence of the following rescuing agents: 100uM Trolox (Sigma, 238813), 50nM ZVAD-FMK (SelleckChem, S7023), 1nM Bafilomycin A1 (Sigma, B1793), 10uM Necrostatin-1s (BioVision, 2263-1), 500nM Ferrostatin-1 (Sigma, SML0583), 1 mM freshly prepared N-acetyl-L-cysteine (Sigma, A9615), 100uM deferoxamine (Sigma, D9533), 5uM IKE (provided by Brent Stockwell). After 48 hours of incubation, 10 μL of AlamarBlue reagent (Thermofisher, 88952) were added to each well containing 100 μL of experimental media. Plates were incubated at standard culture conditions for 4 hours before measuring fluorescent level on a Promega plate reader using the green filter.

For flow cytometric detection of ROS upon cyst(e)inase treatment cells were seeded in triplicates on 6-well plate format at the following density: 100,000 cells per well of PANC-1 and S2-013 cells; 250,000 cells per well of BxPC3 and AsPC-1. Cell were allowed to grow overnight before subjected with 90nM cyst(e)inase treatment in the presence of the same rescuing agents and concentrations, as described above for the cell viability assay. PANC-1, AsPC-1 cells were incubated in the presence of cyst(e)inase and agents for 24 hours; BxPC-3 and S2-013 for 48 hrs. 5uM IKE was added 6-8 hours before the end of the incubation time for each cell lines. After that cells were incubated for 30 minutes in live-cell imaging solution containing 2 μM C-11 BODIPY (Invitrogen). Cells were then washed with PBS, trypsinized with phenol red free 0.5% trypsin (Life Technologies, 25200-056), and neutralized with 10% FBS in PBS at a 1:1 volume. Cells were strained through a 40μM strainer (BD Falcon, 08-771-1), and analyzed a BD Fortessa. A minimum of 10,000 cells were analyzed per condition using FITC channel. Software analysis was carried out using FlowJo V10.

## Mass Spectrometry-Based Metabolomics

### Unlabeled targeted metabolomics (For IKE and PANK inhibitor PD markers)

Cells were plated at 0.5 million cells per well in 6-well plates and treated with the indicated conditions. Following treatment, the medium was aspirated and cells were lysed using cold 80% methanol and extracts were incubated in −80°C for 10 min and centrifugation at 14,000 rpm for 10 min at 4°C. Protein concentration was determined by processing a parallel 6-well plate at equivalent cell density and used to normalize metabolite fractions across samples. Aliquots of the supernatants were then transferred to a fresh microcentrifuge tube and dried. Metabolite extracts were then re-suspended in 35 μl 50:50 MeOH: H2O mixture for LC–MS analysis.

LC-MS analysis was performed using an Agilent Technologies Triple Quad 6470 system ran in negative ion acquisition modes. dMRM transitions and other parameters for each compounds were determined empirically utilizing analytical standards.

Separations were conducted utilizing an Agilent ZORBAX RRHD Extend-C18 column, 2.1 × 150 mm, 1.8 um and ZORBAX Extend Fast Guards. LC gradient profile is: at 0.25 ml/min, 0-2.5 min, 100% A; 7.5 min, 80% A and 20% C; 13 min 55% A and 45% C; 20 min, 1% A and 99% C; 24 min, 1% A and 99% C; 24.05 min, 1% A and 99% D; 27 min, 1% A and 99% D; at 0.8 ml/min, 27.5-31.35 min, 1% A and 99% D; at 0.6 ml/min, 31.50 min, 1% A and 99% D; at 0.4 ml/min, 32.25-39.9 min, 100% A; at 0.25 ml/min, 40 min, 100% A. Column temp is kept at 35 □C, samples are at 4 □C, injection volume is 2 μL.

Mobile phase (A) consists of 97% water and 3% methanol 15 mM acetic acid and 10 mM tributylamine at pH of 5. (C) consists of 15 mM acetic acid and 10 mM tributylamine in methanol. Washing Solvent (D) is acetonitrile. LC system seal washing solvent 90% water and 10% isopropanol, needle wash solvent 75% methanol, 25% water.

Key mass spectrometry parameters utilized were: Gas temp 150 □C, Gas flow 10 l/min, Nebulizer 45 psi, Sheath gas temp 325 □C, Sheath gas flow 12 l/min, Capillary −2000 V, Delta EMV −200 V. Dynamic MRM scan type is used with 0.07 min peak width, acquisition time is 24 min. Delta retention time of plus and minus 1 min, fragmentor of 40 eV and cell accelerator of 5 eV are incorporated in the method.

### Metabolomics Data Analysis (For IKE and PANK inhibitor PD markers)

Raw data were pre-processed with Agilent MassHunter Workstation Software Quantitative QqQ Analysis Software (B.07.00). Metabolite counts were then normalized by the total intensity of all metabolites to reflect equal sample loading. Finally, each metabolite abundance in each sample was divided by the median of all abundance levels across all samples for proper comparisons, statistical analyses, and visualizations among metabolites

### Remaining Metabolomics Data Acquisition and Analysis

For steady state, an Agilent 1290 UHPLC-6490 Triple Quandruple MS system as above was used. For negative ion acquisition, a Waters Acquity UPLC BEH amide column (2.1 x 100mm, 1.7μm) column with the mobile phase (A) consisted of 20 mM ammonium acetate, pH 9.6 in water, and mobile phase (B) was used. Gradient program: mobile phase (B) was held at 85% for 1 min, increased to 65% in 12 min, then to 40% in 15 min and held for 5 min before going to initial condition and held for 10 min. For positive ion acquisition, a Waters Acquity UPLC BEH TSS C18 column (2.1 x 100mm, 1.7μm) column was used with mobile phase A) consisting of 0.5 mM NH4F and 0.1% formic acid in water; mobile phase (B) consisting of 0.1% formic acid in acetonitrile. Gradient program: mobile phase (B) was held at 1% for 1.5 min, increased to 80% in 15 min, then to 99% in 17 min and held for 2 min before going to initial condition and held for 10 min. The column was kept at 40 □C and 3 μl of sample was injected into the LC-MS/MS with a flow rate of 0.2 ml/min. Tuning and calibration of QqQ MS was achieved through Agilent ESI-Low Concentration Tuning Mix.

Optimization was performed on the 6490 QqQ in negative or positive mode individually for each of 220 standard compounds to get the best fragment ion and other MS parameters for each standard. Retention time for each standard of the 220 standards was measured from pure standard solution or a mix standard solution. The LC-MS/MS method was created with dynamic dMRMs with RTs, RT windows and MRMs of all the 220 standard compounds.

In both acquisition modes, key parameters of AJS ESI were: Gas temp 275 □C, Gas Flow 14 l/min, Nebulizer at 20 psi, Sheath Gas Heater 250 □C, Sheath Gas Flow 11 L/min, Capillary 3000 V. For negative mode MS: Delta EMV was 350 V, Cycle Time 500 ms and Cell accelerator voltage was 4 V, whereas for positive acquisition mode MS: Delta EMV was set at 200 V with no change in cycle time and cell accelerator voltage.

The QqQ data pre-processed with Agilent MassHunter Workstation Software Quantitative QqQ Analysis Software (B0700). Additional analyses were post-processed for further quality control in the programming language R. We calculated coefficient of variation (CV) across replicate samples for each metabolite given a cut-off value of peak areas in both the positive and the negative modes. We then compared distributions of CVs for the whole dataset for a set of peak area cut-off values of 0, 1000, 5000, 10000, 15000, 20000, 25000 and 30000 in each mode. A noise cut-off value of peak areas in each mode was chosen by manual inspection of the CV distributions. Each sample is then normalized by the total intensity of all metabolites to reflect the same protein content as a normalization factor. We then retained only those metabolites with at least 2 replicate measurements. The remaining missing value in each condition for each metabolite was filled with the mean value of the other replicate measurements. Finally, each metabolite abundance level in each sample was divided by the median of all abundance levels across all samples for proper comparisons, statistical analyses, and visualizations among metabolites. The statistical significance test was done by a two-tailed t-test with a significance threshold level of 0.05. The p-values were not adjusted in favor of more flexible biological interpretation.

For the ^13^C–cystine/methionine label incorporation studies, an Agilent 1260 UHPLC combined with a 6520 Accurate-Mass Q-TOF LC/MS was utilized. Agilent MassHunter Workstation Software LC/MS Data Acquisition for 6200 series TOF/6500 series QTOF (B.06.01) was used for calibration and data acquisition. A Waters Acquity UPLC BEH amide column (2.1 x 100mm, 1.7μm) column was used with mobile phase (A) consisting of 20 mM NH4OAc in water pH 9.6, and mobile phase (B) consisting of ACN. Gradient program: mobile phase (B) was held at 85% for 1 min, increased to 65% in 12 min, then to 40% in 15 min and held for 5 min before going to initial condition and held for 10 min. The column was at 40 □C and 3 μl of sample was injected into the LC-MS with a flow rate of 0.2 ml/min. Calibration of TOF MS was achieved through Agilent ESI-Low Concentration Tuning Mix.

Key parameters for both acquisition modes were: mass range 100-1200 da, Gas temp 350 □C, Fragmentor 150 V, Skimmer 65 v, Drying Gas 10 l/min, Nebulizer at 20 psi and Vcap 3500 V, Ref Nebulizer at 20 psi. For negative mode the reference ions were at 119.0363 and 980.01637 m/z whereas for positive acquisition mode, reference ions at 121.050873 and 959.9657 m/z

For ^13^C-labeling data analysis, we used Agilent MassHunter Workstation Software Profinder B.08.00 with Batch Targeted Feature Extraction and Batch Isotopologue Extraction and Qualitative Analysis B.07.00. Various parameter combinations, e.g. mass and RT tolerance, were used to find best peaks and signals by manual inspection. Key parameters were: mass tolerance = 20 or 10 ppm and RT tolerance = 1 or 0.5 min. Isotopologue ion thresholds, the anchor ion height threshold was set to 250 counts and the threshold of the sum of ion heights to 500 counts. Coelution correlation threshold was set to 0.3.

All other bioinformatics analyses including graphs and plots were done using R/Bioconductor.

### Laser capture microdissection and RNA sequencing

Cryosections of OCT–embedded tissue blocks from KPFSR mice treated with corn oil and tamoxifen, respectively, were transferred to PEN membrane glass slides and stained with cresyl violet acetate. Adjacent sections were H&E stained for pathology review. Laser capture microdissection was performed on a PALM MicroBeam microscope (Zeiss), collecting at least 10000 cells per sample. Total RNA was extracted using the RNeasy Micro Kit (Qiagen) and amplified using the Clontech SMART-seq v4 Ultra Low Input RNA Kit to create cDNA. Next, the Illumina Nextera XT kit was used to prepare libraries which were then sequenced to a depth of 30 million, 100bp, single-end reads on the Illumina HiSeq4000 platform. Reads were mapped to the UCSC mm10 reference genome and quantified per gene, respectively, using the STAR (v 2.5.2b) and featureCounts (v 1.5.0-p3) software.

### Ferroptosis signatures

First, gene expression data were retrieved from the supplement of Dixon et al.(*25*) as FPKM per gene. Differential gene expression (DEG) analysis was carried out between samples treated with DMSO and erastin, respectively, after log2 transformation using the *limma* R package. Genes significantly upregulated upon erastin treatment at an FDR <= 0.05 (n = 45) represented the *in vitro* ferroptosis signature. Next, DEG analysis was carried out between LCM-RNA-Seq samples retrieved from epithelia of KPFSR tumors whose animals had been treated with corn oil and tamoxifen, respectively, using the *DESeq2* R(*41*) with raw counts as input. Genes significantly upregulated upon tamoxifen treatment at an FDR <= 0.005 (n = 44) represented the *in vivo* ferroptosis signature. Gene set enrichment of the aforementioned signatures on the entire *in vitro* and *in vivo* differential gene expression signature, respectively, was carried out as described previously(*42*).

#### Microarray data

Gene expression data from studies comparing pancreatic ductal adenocarcinoma specimen with normal pancreas parenchyma were downloaded from the Gene Expression Omnibus (GEO) using the *GEOquery* R**(*43*)** package **(*44*)** and the following accession numbers: GSE32676 **(*45*)**, GSE19650 **(*46*)**, GSE16515 **(*47*)**, GSE62452 **(*48*)** and GSE71729 **(*49*)**. U133A-CEL files from **(*50, 51*)** were obtained from ArrayExpress under accession number E-MEXP-950 and normalized using *GCRMA* **(*52*)**. The following samples were excluded because of outlier behavior during exploratory data analysis: NPD15, T55, TPK9, NPK13. For all studies, probes were collapsed at the gene level using their mean normalized expression.

#### TCGA data

RNA-Seq V2 expression, clinical and mutation data were downloaded for all available TCGA tumor types using the *RTCGAToolbox* R package (*53*). The run date was set to “2016-01-28”.

#### ICGC PDA expression data

RNA-Seq expression data and clinical annotation for 96 cases were retrieved from the supplementary data of Bailey et al.(*54*) Illumina HumanHT-12 v4.0 microarray expression data were downloaded from the ICGC data portal for 269 cases. Only those cases were retained for further analyses that were annotated in Bailey et al. and did not have an RNA-Seq expression profile, leaving 142 unique cases.

### Differential gene expression analysis

Genome-wide differential gene expression analysis normal tissue of origin and tumor tissue were calculated using the *limma* (*55*) R package using (gc)rma-normalized expression data for microarray studies and log2-transformed normalized count data for the TCGA cohorts. If normal and tumor samples were matched, i.e. from the same patient, a paired design was specified in the design formula.

### Gene set enrichment analysis

In order to examine the relationship of pathway activity and *SLC7A11* expression, we used the R implementation of single sample Gene Set Enrichment analysis: GSVA (gene set variation analysis) with default parameters (*56*) after pre-filtering each expression matrix for genes with an interquartile range > 0.5. PDA tumors from the following cohorts were examined: TCGA, ICGC RNA-Seq, ICGC microarray, NIH, UNC and Collisson their enrichment score per sample was calculated for the following gene sets from the MSigDB (v.6.0) (*57*): HALLMARK_REACTIVE_OXIGEN_SPECIES, SINGH_NFELE2_TARGETS and GO_CELL_REDOX_HOMEOSTASIS.

### Effect size meta-analysis

#### Log_2_ fold change

The effect size (i.e. log_2_ fold change) and its standard error for *SLC7A11* were extracted from the respective genome-wide differential expression analysis of 6 studies where global expression in normal pancreatic tissue was compared to PDA.

#### Pearson correlation

Pearson correlation and its standard error were calculated for each study and gene set, respectively, between the gene set enrichment scores per sample and the median-centered *SLC7A11* expression per sample using the *cor.test* function from the *stats* R package.

Meta-analysis for each metric was carried out using the *metafor* (*58*) R package. Both random and fixed effect models were fit using the *rma* function (method = “REML” and method = “FE”, respectively).

#### Survival analysis

The association of *SLC7A11* expression status with disease outcome was evaluated using a log-rank test as implemented in the *survdiff* function from the *survival* (*59*) R package. Patients were grouped into tertiles according to normalized *SLC7A11* expression and differences in outcome were assessed between the upper and lower tertile.

### Animal Breeding and genotyping

All studies were carried out in accordance with the relevant institutional guidelines of Columbia University.

### Generation of Pdx1-FlpO mice

The proximal 6kb promoter of the Pdx1-Cre transgene (Gu et al., 2002; Hingorani et al., 2003) was fused to the start codon of mammalian codon-optimized, thermostable Flp recombinase (FlpO) (Raymond and Soriano, 2007), with subsequent fusion of the 5’ end of the FlpO open reading frame to the hGH polyadenylation signal sequence using the In-Fusion cloning system (Clontech) according to the manufacturer’s instructions. The Pdx1-FlpO cassette was excised from a large-scale plasmid preparation, gel-purified, and microinjected into the pronuclei of fertilized FVB oocytes at the Gladstone Transgenic Mouse Core. Following implantation, birth, and weaning, transgene-bearing founder mice were identified via PCR and maintained on an FVB background.

Pdx1-FlpO founders were crossed to homozygous FVB alkaline phosphatase Flp reporter mice (gift, Dr. Susan Dymecki, Harvard University), Pdx1-FlpO-harboring progeny sacrificed at 2 months of age, and skin, brain, liver, pancreas, kidney, lung, stomach, duodenum, small intestine, colon collected and flash frozen in liquid nitrogen. Frozen sections of each were cut, alkaline phosphatase histochemistry performed (Vector Laboratories), and sections examined microscopically. Founder lines exhibiting prominent alkaline phosphatase activity in the pancreas were backcrossed an additional five generations and this process was repeated to identify founder lines exhibiting stable transgene expression.

### Generation of *Slc7a11* conditional knockout mice

Mice bearing an *Slc7a11* conditional null allele (EMMA ID: 10001, strain designation *Slc7a11*^tm1a(EUCOMM)Wtsi^, referred to here as *Slc7a11*^Fl^) were imported from the IMPC repository. These mice were crossed to homozygous ACTB: FLPe mice (Jackson Laboratory, Stock no. 003800) to create *Slc7a11* conditional mice in which loxP sites surround exon three of the gene. Using a combination of the Pdx1-Flpo, KRAS^FSFG12D^ (Jackson Laboratory, Stock no. 008653), p53^R172H^ (Jackson Laboratory, Stock no. 008652), Rosa26^CreERT2^ (Jackson Laboratory, Stock no. 008463), and our newly made *Slc7a11*^Fl/Fl^ mice, we were able to generate a new strain of genetically engineered mouse model akin to the *KRAS*^*LSL-G12D*/+^;*p53^LSL-R172H^;Pdx1-Cre* (KPC) mouse (*21*), but in which we have temporal control of the deletion of *Slc7all* using tamoxifen. We term these mice *KRAS*^*FSF-G12D*/+^*; p53^R172H^; Pdx1-Flpo^tg/+^; Slc7a11^Fl/Fl^; Rosa26^CreERT2/+^*, KPFSR mice.

### Genotyping

Genotyping was carried out using protocols provided by Jackson Laboratory for all strains obtained from this vendor. *Slc7a11^Fl/Fl^* genotyping was carried out as follows:

Forward primer: 5’-tgggttggtctctggtgatc-3’; Reverse primer: 5’-cctgtgaagatccgcctact-3’
Cycling conditions: 1. 94°C 3 minutes; 2. 94°C 1 minute; 3. 60°C 2 minutes; 4. 72°C 1 minute; Cycle to step 2, 29 times.; 5. 72°C 5 minutes; 6. 4°C forever
Expected amplicons: *Slc7a11*^Fl/Fl^ mice: 386 bp; WT mice: no amplicon *Slc7a11* recombinatorial genotyping was custom designed as follows:
Forward primer: 5’-tgg gtt ggt ctc tgg tga tc-3’; Reverse primer: 5’-ctt aac ccc agc acc att cg-3’
Cycling conditions: 1. 95°C 2 min; 2. 95°C 1 min; 3. 56°C 30 seconds; 4. 72°C 1 min; Cycle to step 2, 34 times; 5. 72°C 5 min; 6. 4°C forever
Expected amplicons: *Slc7a11*^Fl/Fl^ unrecombined = 1285 bp; *Slc7a11*^Fl/Fl^ recombined = 450 bp; WT mouse = 1240 bp.

### Western blot

Western blot for Slc7a11 was carried out using standard protocols. In short, cultured MEFs were collected in PBS and lysed with Flag lysis buffer. Tumor samples were ground and lysed in Flag lysis buffer, and proteins were quantified with Bradford assay. Proteins were run on an SDS PAGE gel under denaturing conditions, transferred to a membrane, and probed with primary antibody (Cell Signaling, s12691) overnight at 4°C. Blot was probed with an appropriate secondary and developed using a standard ECL kit.

### Survival study

At approximately 42 days of age, KPSFR mice were treated with cerulein (Sigma, C9026) at 250 μg/kg by intraperitoneal injection for 5 consecutive days to induce chronic pancreatitis and accelerate tumor formation, as per previous reports(*60–62*). Tumor formation was monitored initially by once weekly palpation and upon positive palpation, by twice-weekly ultrasound. Once the average diameter of tumors of the pancreas was 4-7 mm in size, mice were randomly enrolled into the survival study. Mice were treated with either corn oil (Sigma, C8267) or tamoxifen (Sigma, T5648) at 200 mg/kg by oral gavage for 6 consecutive days. Afterwards, a small cohort of mice was enrolled on a tamoxifen and NAC combination treatment. These mice were administered tamoxifen in the aforementioned fashion and administered NAC in their drinking water at 1 g/L. Subsequently, small cohorts of KPC mice were enrolled in a vehicle treatment arm and on an NAC only treatment arm, as controls. Mice were euthanized once they reached endpoint criteria consisting of a combined physical and behavioral metric designed to identify mice prior to death. Tumor samples were either fixed in formalin overnight at 4°C, fixed in PFA at 4°C overnight followed by sucrose-mediated water displacement for 24 hours at 4°C, frozen in OCT, or flash frozen in liquid nitrogen.

### Ultrasound

Tumor ultrasonography and volume quantification was carried out as previously described(*63*).

### Pathology and immunohistochemistry

Samples that had been fixed in formalin were washed in 70% ethanol and subjected to standard dehydration processing to prepare them for mounting in paraffin wax blocks. Paraffin blocks were sectioned on a Leica RM 2235 at 5 μM thickness. The sections were mounted on charged slides and heated to 60°C to melt wax and ensure tissue adherence to the slide. Slides were then subjected to standard rehydration and antigen retrieval was carried out for five minutes in boiling 10 mM sodium citrate buffer pH 6, .05% Tween-20 using a pressure cooker. Slides were allowed to cool to room temperature in an ice-cold water bath. Slides were then incubated in 3% hydrogen peroxide for 20 minutes at room temperature to block endogenous peroxidases. Slides were blocked in 1.5% horse serum and 2% animal free blocker (Vector Laboratories, SP-5030) for 1 hour at room temperature. Malondialdehyde-stained slides however were blocked in 5% BSA. Slides were then stained with the appropriate antibody: Cleaved caspase-3 (Cell Signaling, 9664S), 1:100 dilution; Ki67 (Cell Signaling, 12202S), 1:100 dilution; phosphohistone-H3 (Cell Signaling, 9716S), 1:100 dilution; 4-hydroxynonenal (Abcam, 46545), 1:200. Primary antibody incubation was carried out overnight at 4°C. Slides were then washed 3x with PBS-T and incubated with the appropriate secondary antibody for 30 minutes at room temperature. Staining was developed using the DAB reagent (Vector Labs, VV-93951085). Quantification for CC3, Ki67, and pHH3 was done in a blinded fashion, counting positive cells per 40x field for 10 fields per sample. Quantification for 4HN staining was done by first deconvolving hematoxylin and DAB staining using Fiji. The DAB image component was then adjusted for a threshold of 175 to identify positive staining. Finally, Fiji was used to analyze particle number using a circularity of 0-1 and a size above 200 pixels, yielding a total particle count per low powered (1.25X) field per sample.

Histological staining on paraffin sections (hematoxylin and eosin, Oil Red O) were carried out using standard protocols. Following digital capture on Olympus BX51, histology images were processed using Affinity Photo software, using three filters/adjustments applied evenly across the entire image: unsharp mask, white balance, and levels adjustment.

### Transmission Electron Microscopy

Tissues were fixed in a solution of 2% paraformaldehyde, 2.5% glutaraldehyde, and 2mM CaCl2 in 0.15 M sodium cacodylate buffer (pH 7.4) for 2 hours at room temperature. They were then post-fixed in 1% osmium tetroxide for 40 minutes and 1.5% potassium ferricyanide in sodium cacodylate buffer for 1 hour at room temperature in the dark. Tissues were stained en bloc in 1% aqueous uranyl acetate (4°C in the dark) for 1 hour, dehydrated in a series of graded acetones, and embedded in Eponate12 resin (Ted Pella). Ultra-thin sections (70 nm) were obtained using a diamond knife (Diatome) in an ultramicrotome (Leica EM UC7) and placed on copper grids (300 mesh). Sections were imaged on a Zeiss Libra 120 TEM operated at 120 kV using Zeemas acquisition system with 2K of resolution.

## Supplementary Information

Supplementary videos and tables are available in the online version of the paper.

## Data Availability Statement

The RNA seq datasets generated during this study are available: GEO Accession number: GSE119628; Reviewer token: mruxoimavpgllkd.

## Acknowledgements

We are grateful to fellow members of the Olive, Lyssiotis, and Stockwell labs for technical advice, helpful discussions, and thoughtful critiques. We also thank S.W. Novak for advice on electron microscopy analysis and interpretation. This research was funded in part through the NIH/NCI Cancer Center Support Grant P30CA013696, and used the Confocal and Specialized Microscopy, Molecular Pathology, Flow Cytometry Core, and OPTIC shared resources. M.A.B was supported by a training grant (T32 A009503) and a pre-doctoral fellowship (F31 CA180738). K.P.O received support for this work from the Lustgarten Foundation for Pancreatic Cancer Research (2011 Innovator Award) and the NIH/NCI (1R01CA215607). C.A.L was supported by a Pancreatic Cancer Action Network/AACR Pathway to Leadership award (13-70-25-LYSS); Dale F. Frey Award for Breakthrough Scientists from the Damon Runyon Cancer Research Foundation (DFS-09-14); Junior Scholar Award from The V Foundation for Cancer Research (V2016-009); and Kimmel Scholar Award from the Sidney Kimmel Foundation for Cancer Research (SKF-16-005). V.P. was supported by a PRCRP Horizon award by the Department of Defense (W81XWH-17-1-0497). Metabolomics studies performed at the University of Michigan were supported by NIH grant U24-DK097153. B.R.S. is supported by the NCI/NIH (R35CA209896 and P01CA087497).

## Author Contributions

Conceptualization, M.A.B. and K.P.O; Methodology, M.A.B., T.H., E.S.S., C.A.L., B.R.S., W.G., U.M., L.R.A., G.M.W., K.P.O.; Investigation, M.A.B., H.L., H.M.C., D.K., V.P., P.S., C.M.F, L.Z., Z.P.T., S.A.S., C.F.P., K.E.D., A.D., T.L., A.I. T.H.; Resources, E.S.S., B.R.S., C.A.L., C.L., A.M., J.K., A.D., M.A.B., K.P.O.; Writing – Original Draft, M.A.B. and K.P.O.; Writing – Review and Editing, M.A.B., H.C.M., V.P., H.L., A.D., C.F.P., C.A.L., B.R.S., K.P.O.; Visualization, M.A.B., C.M.F., H.L., H.C.M., E.S.S., C.A.L., K.P.O.; Supervision, G.M.W., B.R.S., C.A.L., K.P.O.; Funding Acquisition, M.A.B., C.A.L., B.R.S., and K.P.O.

## Author Information

B. Stockwell holds equity in and serves as a consultant to Inzen Therapeutics, and is an inventor on patents and patent applications related to IKE and ferroptosis. G. Georgiou and E. Stone are inventors on intellectual property related to this work and have an equity interest in Aeglea Biotherapeutics, a company that has licensed the commercial development of Cyst(e)inase. The other authors declare no competing financial interests. Correspondence and requests for materials should be addressed to K.P.O.

