## Supplementary Figure 1 for "“Induction of pancreatic tumor-selective ferroptosis through modulation of cystine import”"

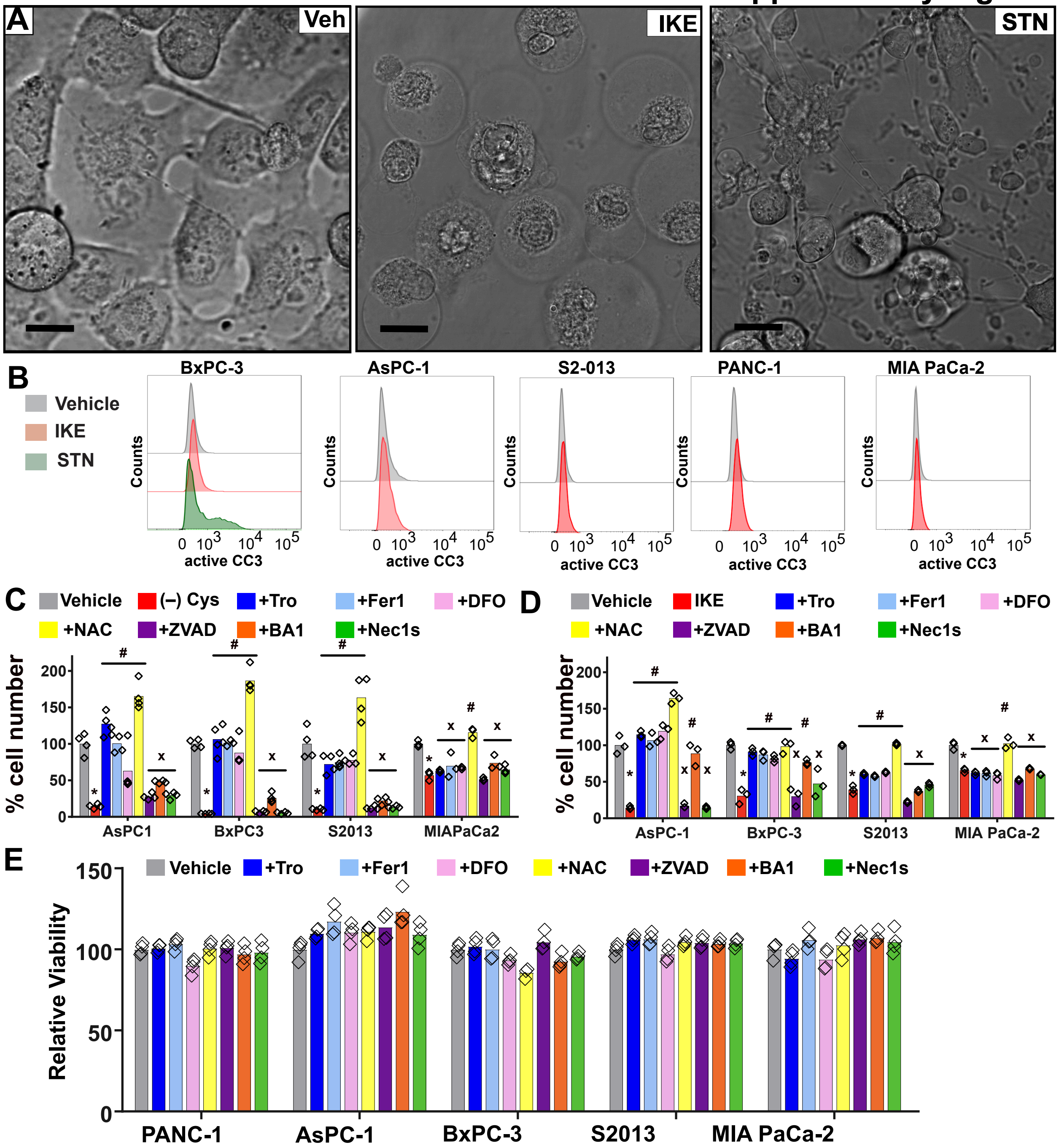


**Supplementary Figure 1: The mechanism of ferroptosis in PDA cell lines (A)** High magnification image of PANC-1 cells cultured in the present of vehicle (0.1% DMSO), 5 µM IKE or 0.2 µM staurosporine (STN), a known inducer of apoptosis, after 16 hours. Bar = 15 µm. **(B)** Flow cytometric analysis of cleaved caspase 3 activation in cell lines stained with a FITC-based active caspase 3 antibody, treated with vehicle (0.1% DMSO, gray) and IKE (5 uM, red). In BxPC-3 cells, staurosoporine (STN) treatment is used as a positive control (0.2uM, green). **(C)** Cell viability of a panel of pancreatic cancer cells cultured in vehicle (0.1% DMSO and 1mM HCl veh, shown in gray), the absence of cystine (no cys, shown in red), and in the absence of cystine but in the presence of 100 µM Trolox (Tro, shown in blue), 500 nM ferrostatin-1 (Fer-1, ferroptosis inhibitor, shown in light blue), 100 µM deferoxamine (DFO, iron chelator, shown in pink), 1 mM NAC (NAC, shown in yellow), 50 µM ZVAD-FMK (ZVADFMK, apoptosis inhibitor, shown in purple), 1 nM bafilomycin A1 (BA1, autophagy inhibitor, shown in orange), and 10 µM Necrostatin 1s (Nec1s, necroptosis inhibitor, shown in green). Viability was assessed after 24 hours of treatment. Error equals +/- SEM. N = 3 biological replicates. * = $p < .05$ by a one-way ANOVA followed by a post-hoc Tukey test, compared to vehicle. # = $p < .05$ by a one-way ANOVA followed by a post-hoc Tukey test, compared to (-) cys. x = no statistically significant difference compared to (-) cys. **(D)** Cell viability of a panel of pancreatic cancer cells cultured in vehicle (0.2% DMSO, veh, shown in gray), 5 µM IKE (shown in red), and the aforementioned compounds and the concentrations indicated in Supplemental Figure 1C. Viability was assessed at 24 hours after treatment. Error equals +/- SEM. N = 3 biological replicates. * = $p < .05$ by a one-way ANOVA followed by a post-hoc Tukey test, compared to vehicle. # = $p < .05$ by a one-way ANOVA followed by a post-hoc Tukey test, compared to IKE. x = no statistically significant difference compared to IKE.**(E)** Single treatment controls for all experiments shown in Figure 1C, D; Supplementary Figure 1C, D.
