## Supplementary Figure 2 for "“Induction of pancreatic tumor-selective ferroptosis through modulation of cystine import”"

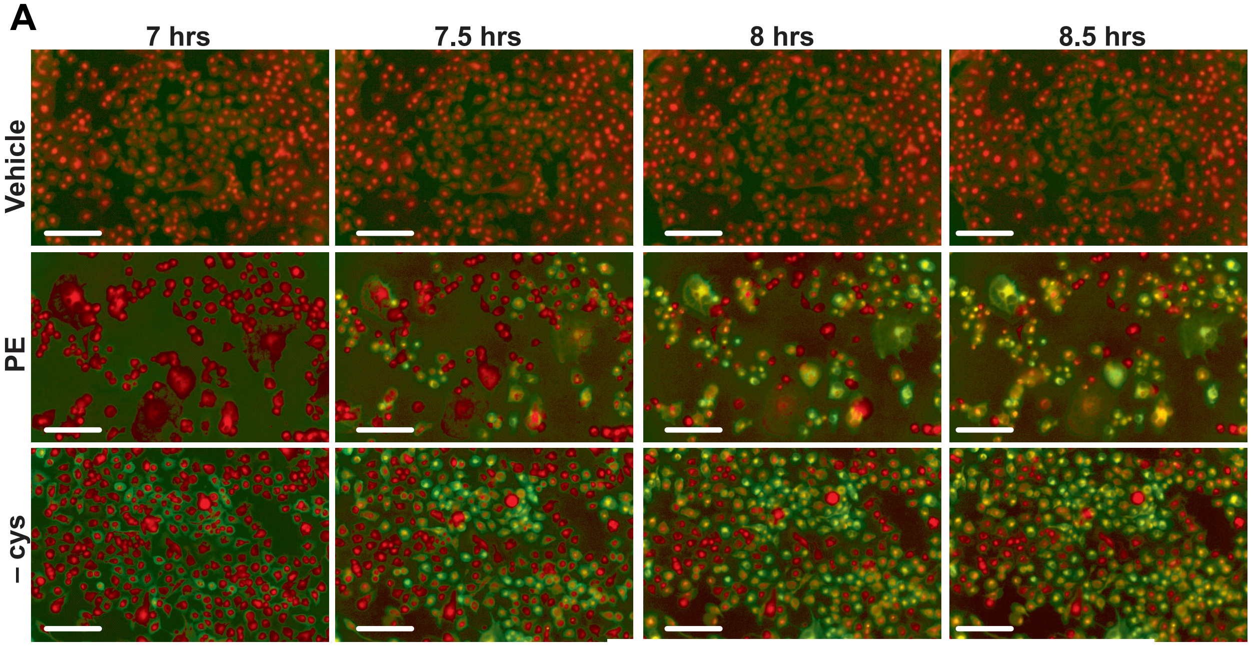


**Supplementary Figure 2: Lipid oxidation in PDA cell lines followign cysteine depletion. (A)** Time-lapse fluorescent images of PANC-1 cells cultured in indicated conditions for indicated times. - cys indicates no cystine present in extracellular media. PE was present at 1 μM. Here, vehicle is 0.01% DMSO. Cells are stained with C-11 BODIPY, a lipid ROS indicator. Green staining highlights oxidized lipids and red staining shows reduced lipids. White bars indicate scale  of 300 μm.
