## Supplementary Figure 3 for "“Induction of pancreatic tumor-selective ferroptosis through modulation of cystine import”"

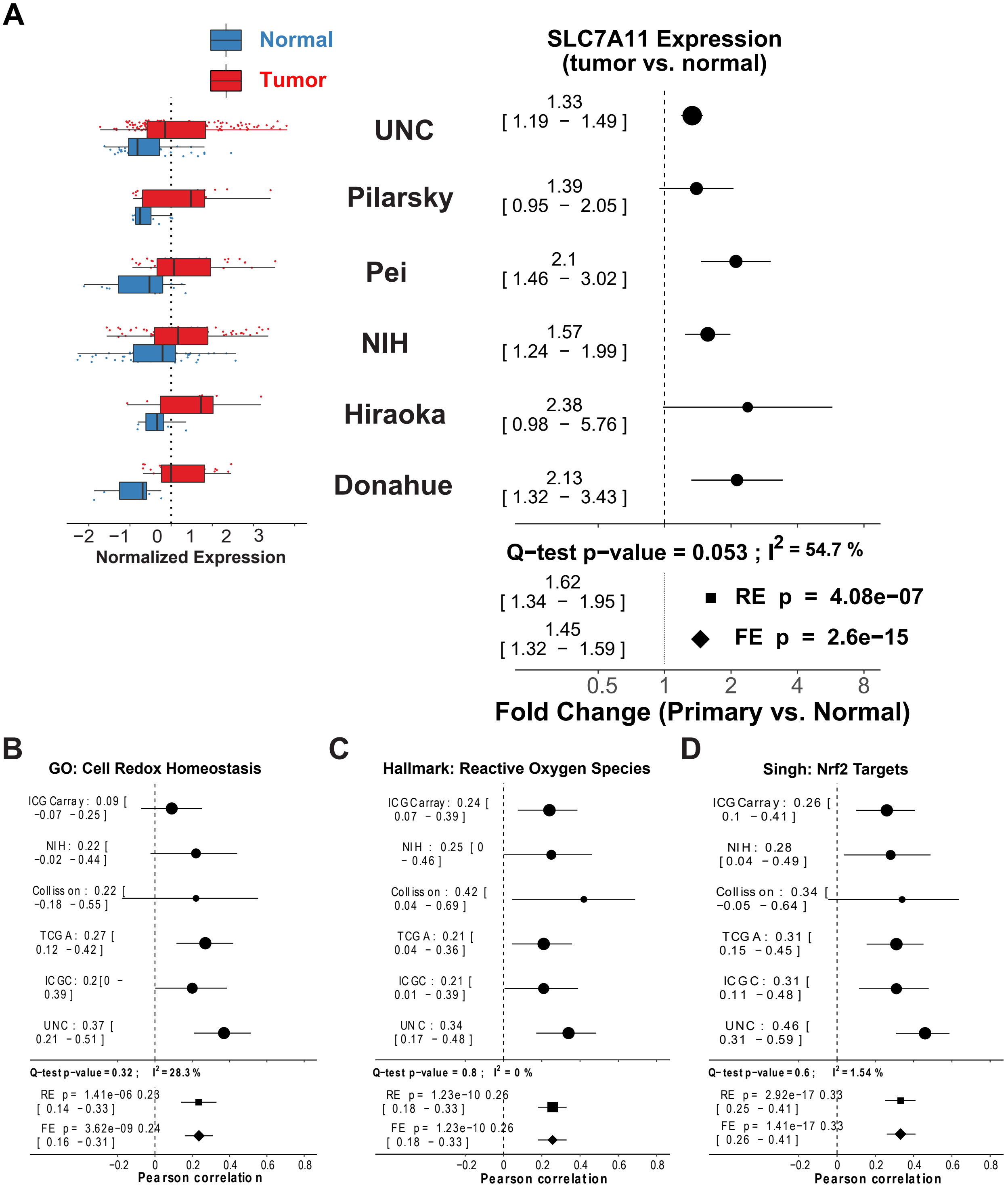


**Supplementary Figure 3. Expression and outcome associations of SLC7A11. (A)** SLC7A11 is overexpressed in primary PDA tumors (red) as compared to normal control (blue) tissue in multiple pancreatic cancer data sets. Effect size estimates of data are shown in the right half of the panel with their 95% confidence interval per study as well as meta-analytic summaries using random (RE) and fixed effect (FE) models. Q-test and $I^2$ indicate measures of inter-study heterogeneity. **(B-D)** Correlation of SLC7A11 expression with curated gene sets. Panels show effect size estimates with their 95% confidence interval per study as well as meta-analytic summaries using random (RE) and fixed effect (FE) models. Q-test and $I^2$ indicate measures of inter-study heterogeneity.
