## Supplementary Figure 4 for "“Induction of pancreatic tumor-selective ferroptosis through modulation of cystine import”"

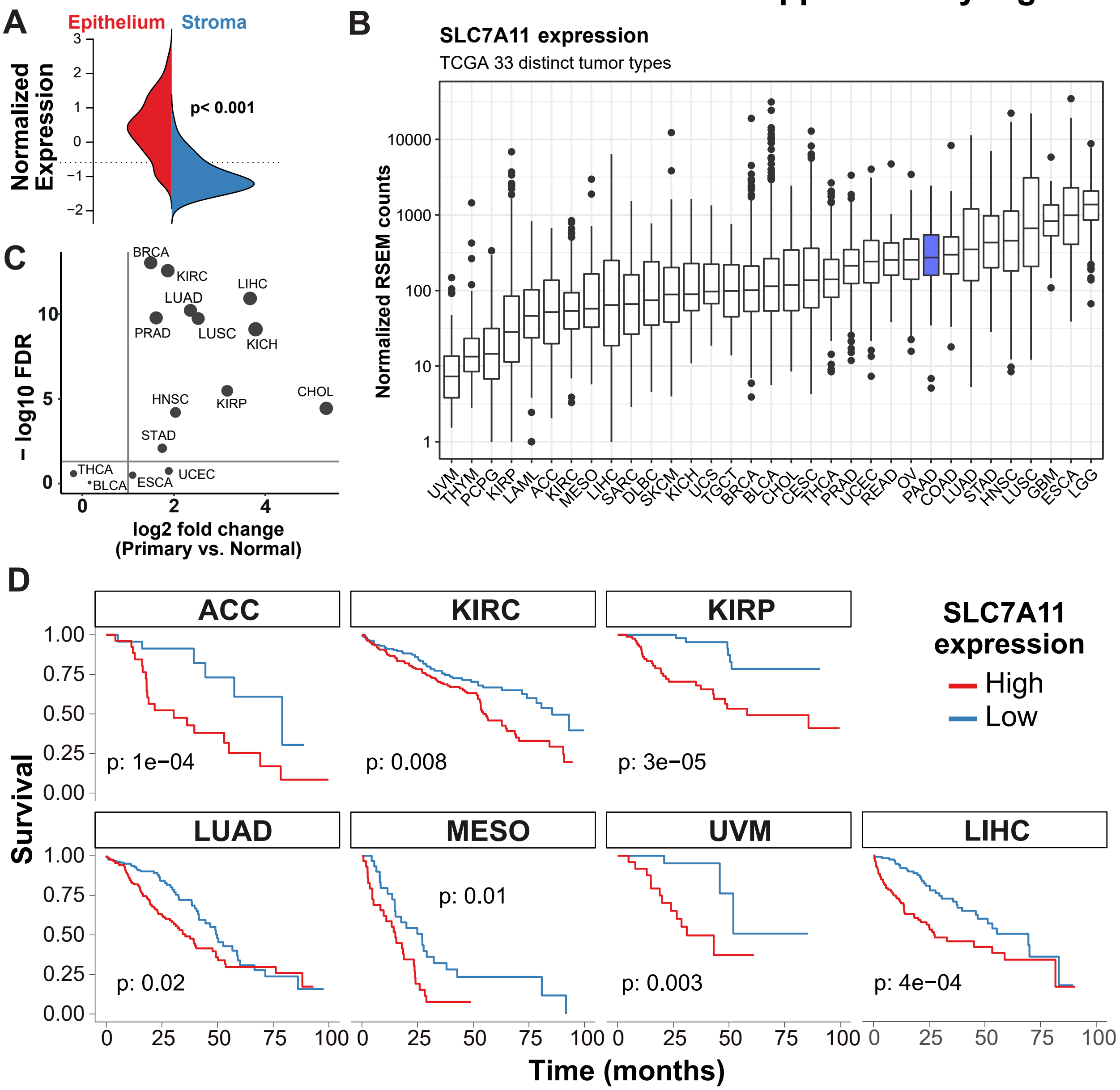


**Supplementary Figure 4. Expression and protein interactions of SLC7A11 (A)** LCM RNA-Seq from matched human PDA epithelium and stroma shows enrichment of expression in epithelial tissue (distribution shown in red) when compared to neighboring stromal tissue, which is not malignant (distribution shown in blue). **(B)** Normalized expression of SLC7A11 across 31 tumor types from TCGA data showing high overall expression in pancreatic ductal adenocarcinoma (PAAD, blue), with generally low variance. **(C)** Analysis of SLC7A11 mRNA expression in TCGA data sets of tumors for which at least 6 normal samples were available shows overexpression of SLC7A11 in most tumor types. **(D)** In TCGA tumors for which there is a difference in outcome between patients expressing varying levels of SLC7A11, high levels are consistently associated with a worse prognosis (logrank test p-values indicated, survival curves shown in red). Abbreviations used according to standard TCGA nomenclature.
