## Supplementary Figure 5 for "“Induction of pancreatic tumor-selective ferroptosis through modulation of cystine import”"

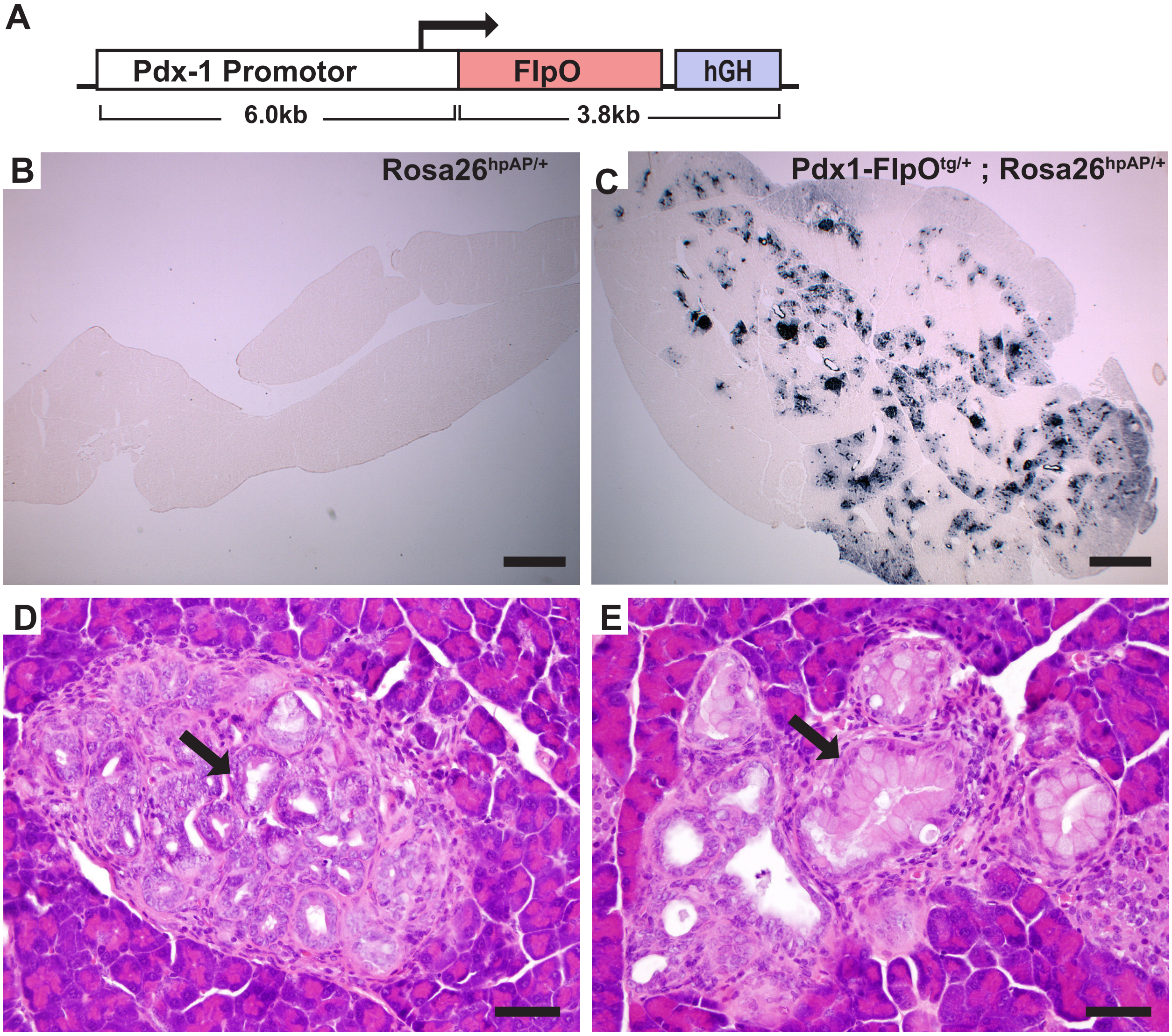


**F**

| KPFSR tissues | Pancreas | | Non-pancreas | |
|---|---|---|---|---|
| | Vehicle | Tamoxifen | Vehicle | Tamoxifen |
| Effective Genotypes: | $Kras^{12D/+}$ | $Kras^{12D/+}$ | $Kras^{+/-}$ | $Kras^{+/-}$ |
| | $p53^{172H/+}$ | $p53^{172H/+}$ | $p53^{+/-}$ | $p53^{+/-}$ |
| | $Slc7a11^{+/+}$ | $Slc7a11^{-/-}$ | $Slc7a11^{+/+}$ | $Slc7a11^{-/-}$ |

**Supplementary Figure 5. Pdx1-FlpO allele design and validation.** (**A**) Design of the Pdx1-FlpO allele. (**B,C**) Pdx1-FlpO founders were crossed to alkaline phosphatase Flp reporter mice (gift, Dr. Susan Dymecki, Harvard University) to visualize recombination in the pancreas. Frozen sections of pancreata from $Rosa26^{hpAP/+}$ (B, negative control) or Pdx1-FlpO; $Rosa26^{hpAP/+}$ (C). Mice were stained for alkaline phosphatase activity (dark blue). Founder lines exhibiting prominent alkaline phosphatase activity in the pancreas were used in further breeding. Bars = 200 µm. (**D,E**) The Pdx1-FlpO strain was crossed with additional strains to generate $Kras^{LSL.G12D/+}$; $p53^{R172H/+}$; $Pdx1\text{-}FlpO^{tg/+}$; $Slc7a11^{Fl/Fl}$ (KPFS) mice. Histopathological examination of the pancreas of young KPFS mice revealed the spontaneous formation of both acinar-to-ductal metaplasia (ADM indicated with arrow, panel D) and pancreatic intraepithelial neoplasia (PanIN indicated with arrow, panel E), both precursors to tumor development. Bar = 50 µm. (**F**) Table indicating effective genotypes of tissues in the KPFSR mouse.
