## Supplementary Figure 6 for "“Induction of pancreatic tumor-selective ferroptosis through modulation of cystine import”"

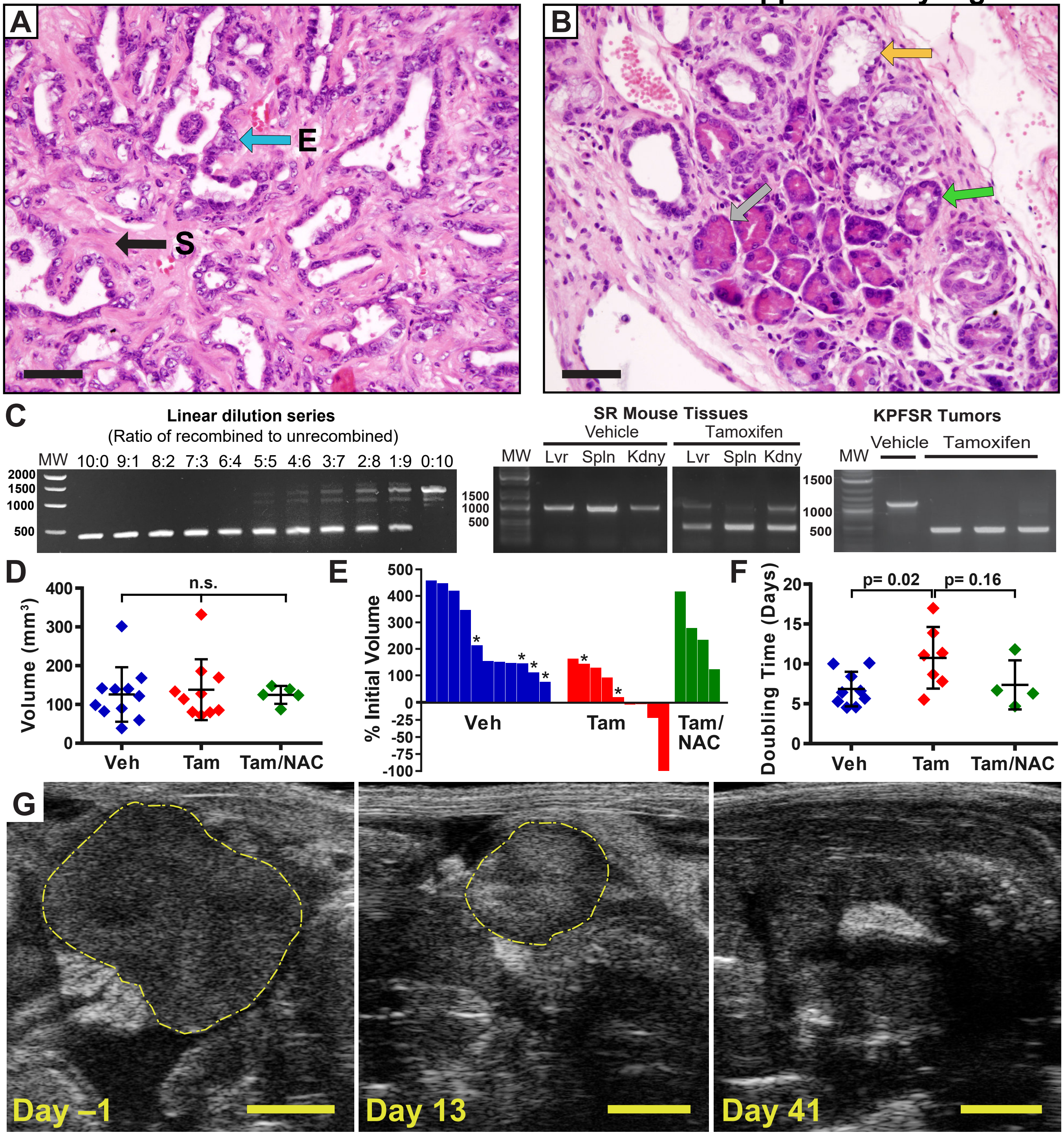


**Supplementary Data Figure 6. Analysis of KPFSR tumors. (A)** Representative KPFSR tumor stained with hematoxylin and eosin (H&E). Desmoplastic stroma is stained in pink while nuclei are stained in purple. Scale bar is 50 μm. Blue arrow indicates malignant epithelial structure (E) while the black arrow indicates the stromal component (S). **(B)** Representative precursor lesions from KPFSR pancreas stained with H&E. Gray arrow indicates a normal acinus, green arrow indicates acinar-to-ductal metaplasia, and yellow arrow indicates an early PanIN lesion. Scale bar is 50 μm. **(C)** Left panel: Dilution series of mixtures of completely recombined DNA to unrecombined DNA, in the indicated ratios. Note preferential detection of recombined allele, with detection of only a faint unrecombined band in the 4:6 lane. Middle panel: recombination as detected by PCR in KPFSR tumors treated for 6 days. Right panel: Analysis of DNA recombination in tissues from SR mice treated with tamoxifen by PCR. Unrecombined, 1285bp; Recombined, 450bp. **(D)** Analysis of tumor volumes at the time of enrollment. Not statisticially significant by one-way ANOVA and posthoc Tukey test. **(E)** Waterfall plots of tumor growth that shows either best regression relative to day 0 or the % tumor volume increase at day 10 (interpolated value for all tumors lacking an ultrasound at day 10. Mice that died prior to day 10 are indicated by an asterisk. **(F)** Tumor growth rates from tumors in the survival study. Error = +/- SD, n = 9 for vehicle, n = 7 for tamoxifen, n = 4 for tamoxifen/NAC (only tumors with at least 4 volumes are utilized in analysis). Statistical analysis carried out by one-way ANOVA followed by posthoc Tukey test. **(G)** Ultrasounds from a tamoxifen-treated KPFSR mouse that underwent complete regression, on study days -1, 13, and 41 of treatment. Hashed yellow lines indicates tumor. Scale bar = 2 mm.
