## Supplementary Figure 7 for "“Induction of pancreatic tumor-selective ferroptosis through modulation of cystine import”"

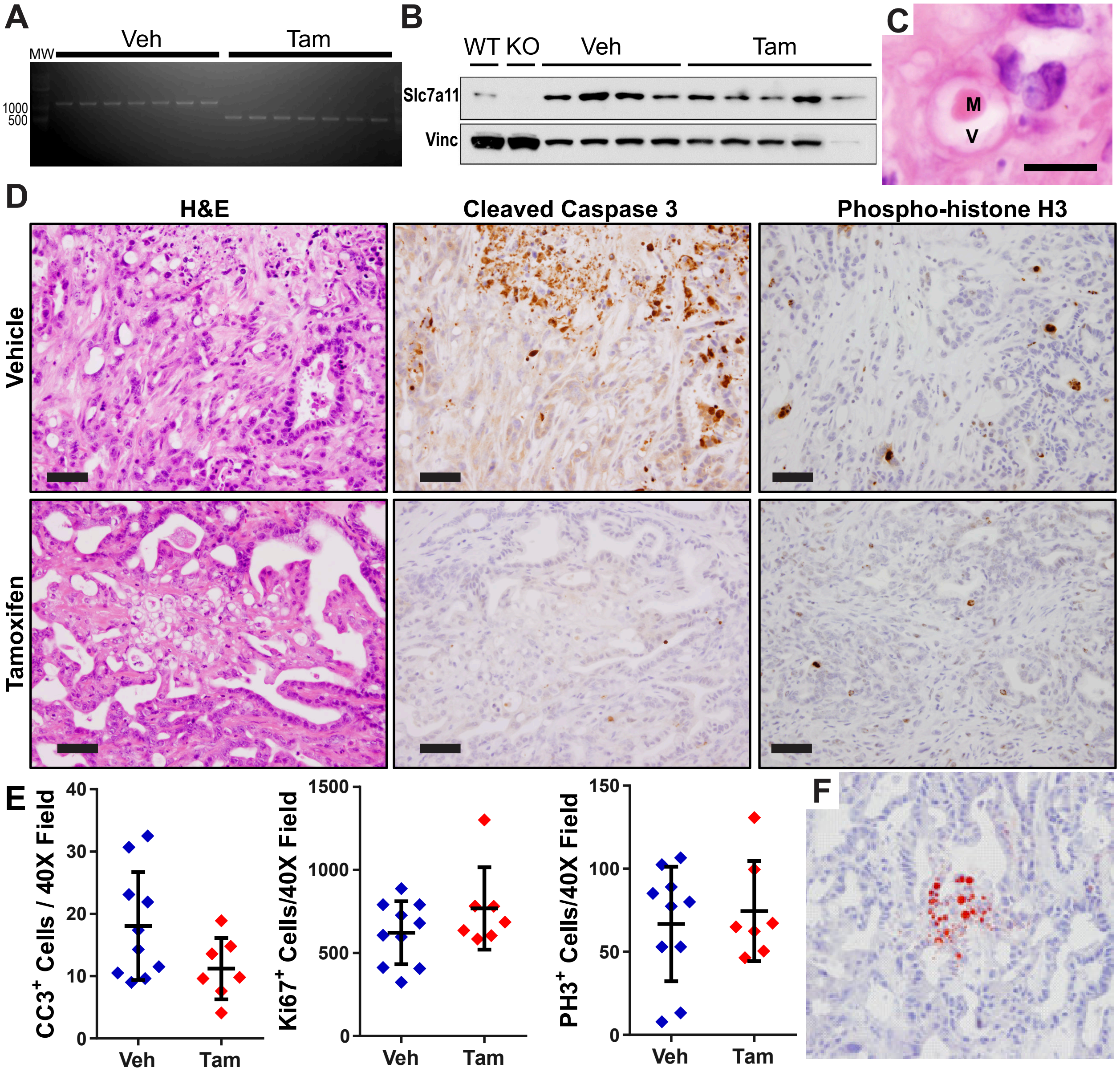


**Supplemental Figure 7: Histopathological description of lesions in tamoxifen-treated KPFSR tumors. (A)** PCR analysis of recombination in endpoint tumors from randomized survival study. Results indicate > 60% recombination of Slc7a11 in endpoint samples. **(B)** Western blot of Slc7a11 in endpoint tumor samples show limited changes in protein levels in control or tamoxifen treated groups. Controls are WT mouse embryonic fibroblasts (MEFs) and knockout (MEFs) generated from our conditional Slc7a11 allele. **(C)** High magnification image of novel lesion, containing megamitochondria (M) and cytoplasmic vacuolization (V). Scale bar = 10 μm. **(D)** Hematoxylin and eosin (H&E), cleaved caspase 3 (CC3) and phosphohistone h3 (pHH3) staining of KPFSR tumors from vehicle and tamoxifen treated tumors. Tamoxifen treated sample shows an example of tissue damaged from Slc7a11 deletion. This region appears CC3 negative but does not exhibit notable changes in pHH3 staining. Scale bar indicates 50μm **(E)** Quantification of CC3, Ki67, and pHH3 staining from both treatment groups. No statistically significat changes are detected by Student's t test when comparing vehicle and tamoxifen groups for any stain. **(F)** Oil Red O staining of tamoxifen treated tumors showing lipid droplets (in red) of unusual size. Scale bar is 50 μm.
