## Supplementary Figure 8 for "“Induction of pancreatic tumor-selective ferroptosis through modulation of cystine import”"

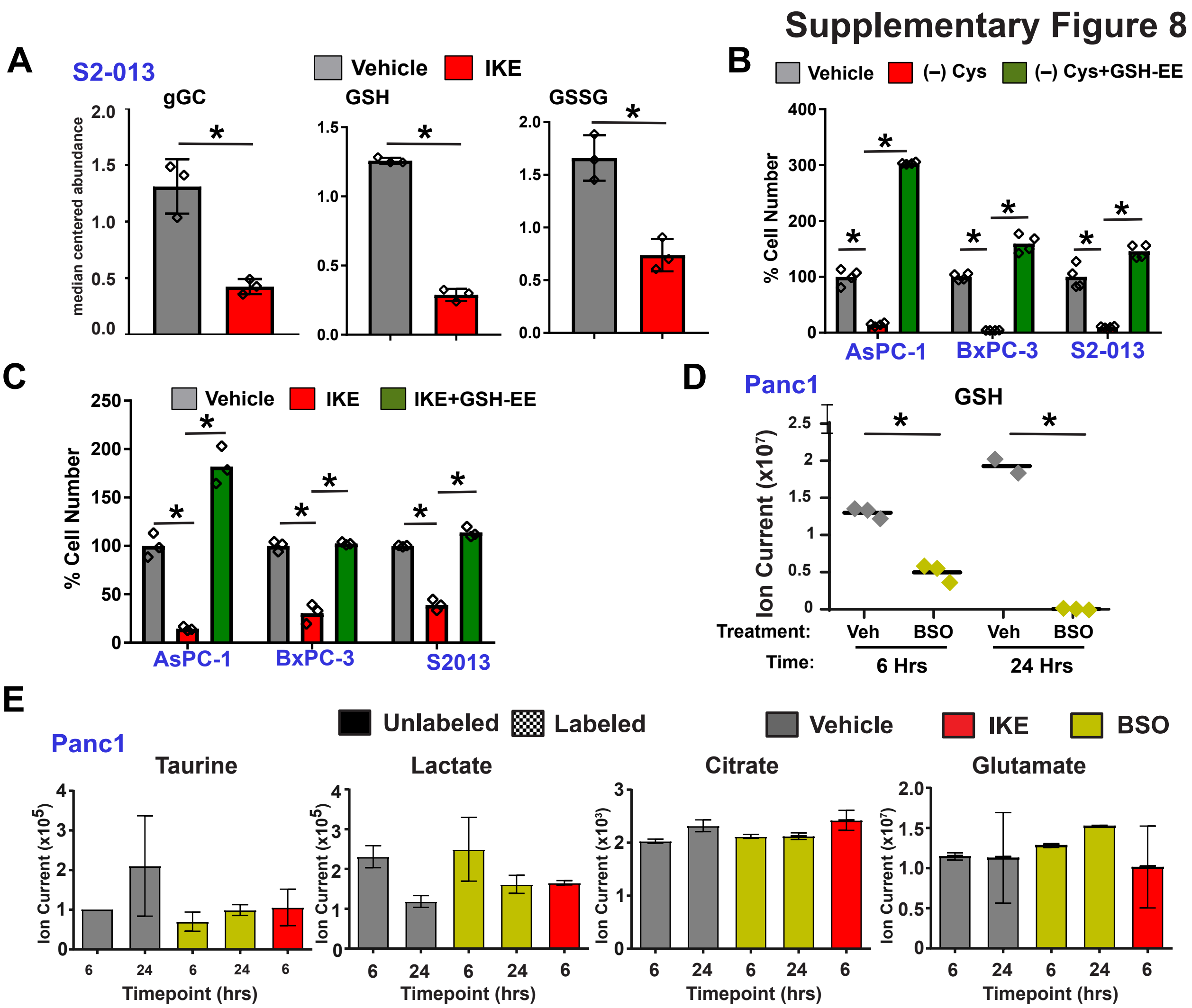


**Supplementary Figure 8. Metabolomics analysis of cysteine utilization and rescue of cell death by cell permeable GSH. (A)** Measurement of gamma glutamyl cysteine (gGC), GSH, and GSSG in S2-013 cells treated with 0.1% DMSO (veh, red) or 5uM IKE (IKE, red) for six hours. * indicated statistical significance by Student's t-test. **(B,C)** GSH-EE rescue of cystine withdrawal (B) and system $x_C^-$ inhibition in sensitive cell lines. * indicates statistical significance ($p < .05$) by one-way ANOVA and posthoc Tukey test. **(D)** GSH levels as measured by mass spectrometry in PANC-1 cells treated with vehicle or BSO for listed timepoints. * indicates statistical significance by Student's t test. **(E)** Measurement of labeled and unlabeled levels of taurine, lactate, glutamate, and citrate in PANC-1 treated with vehicle (.05% DMSO), 5 μM IKE, or 600 μM BSO for indicated times. (All comparisons not significant by one way ANOVA and posthoc Tukey test). No labeled species for these metabolites were detected.
