## Supplementary Figure 9 for "“Induction of pancreatic tumor-selective ferroptosis through modulation of cystine import”"

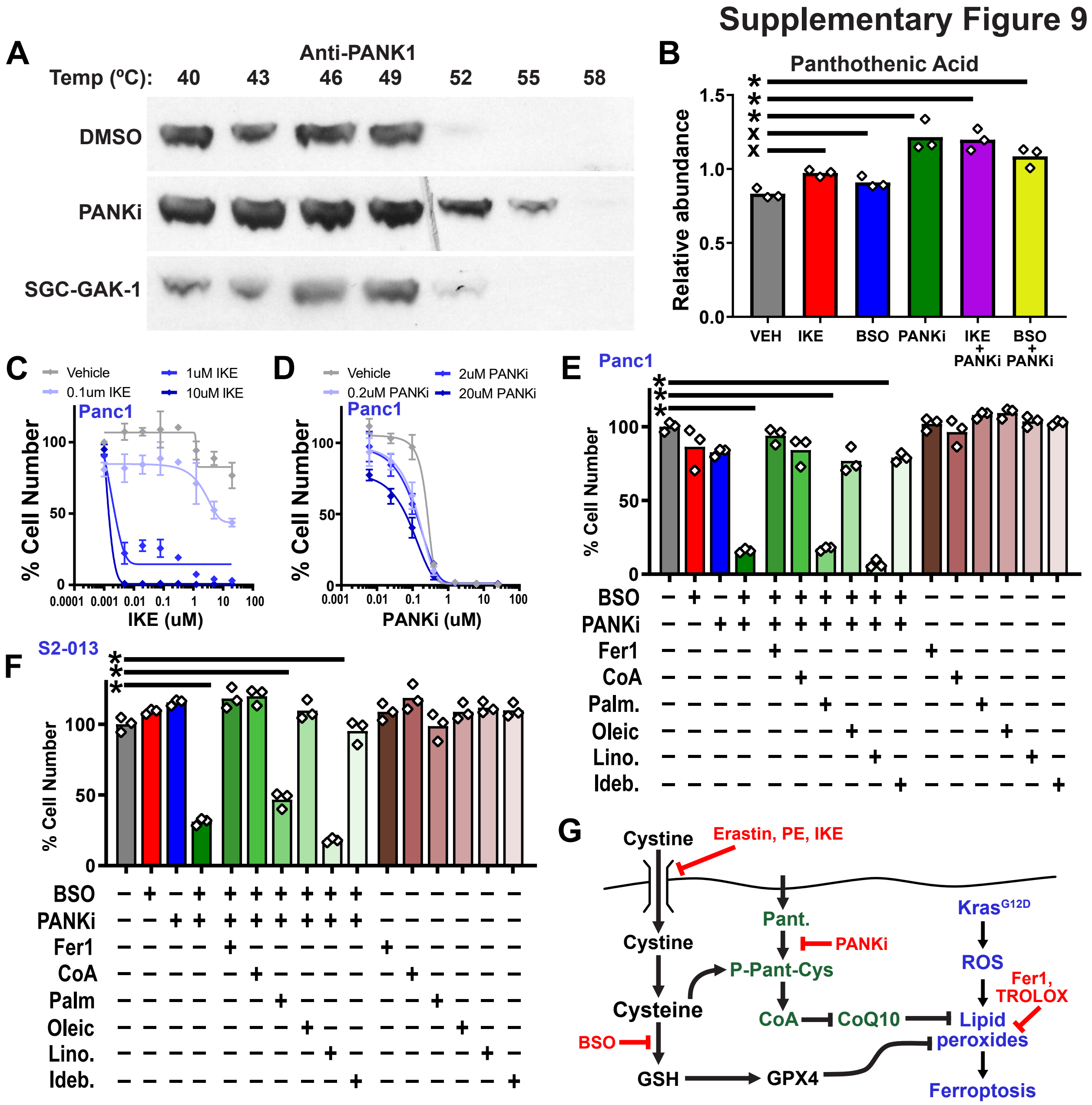


**Supplemental Figure 9. Assessment of combined effects of BSO and PANKi (A)** Cell based thermal shift assay confirms binding of PANK1 inhibitor to target. **(B)** Mass spectrometry measurements of pantothenic acid in given treatment conditions: vehicle (0.1% DMSO), IKE (5uM), BSO (75uM), iPANK (5uM), and various combinations. * indicates statistical significance ($p < .05$) by one-way ANOVA and posthoc Tukey test. **(C)** Dose response curves of PANKi in combination with different concentrations of IKE. **(D)** Dose response curves of IKE with various concentrations of PANKi. **(E, F)** Treatment of two PDA cell lines with combinations of BSO, PANKi, Fer-1, CoA, palmitic acid, oleic acid, linoleic acid, and idebenone. * indicates statistical significance ($p < .05$) by one-way ANOVA and posthoc Tukey test. **(G)** Model of the roles of cysteine utilization in suppressing ferroptosis.
