## Supplementary Figure 10 for "“Induction of pancreatic tumor-selective ferroptosis through modulation of cystine import”"

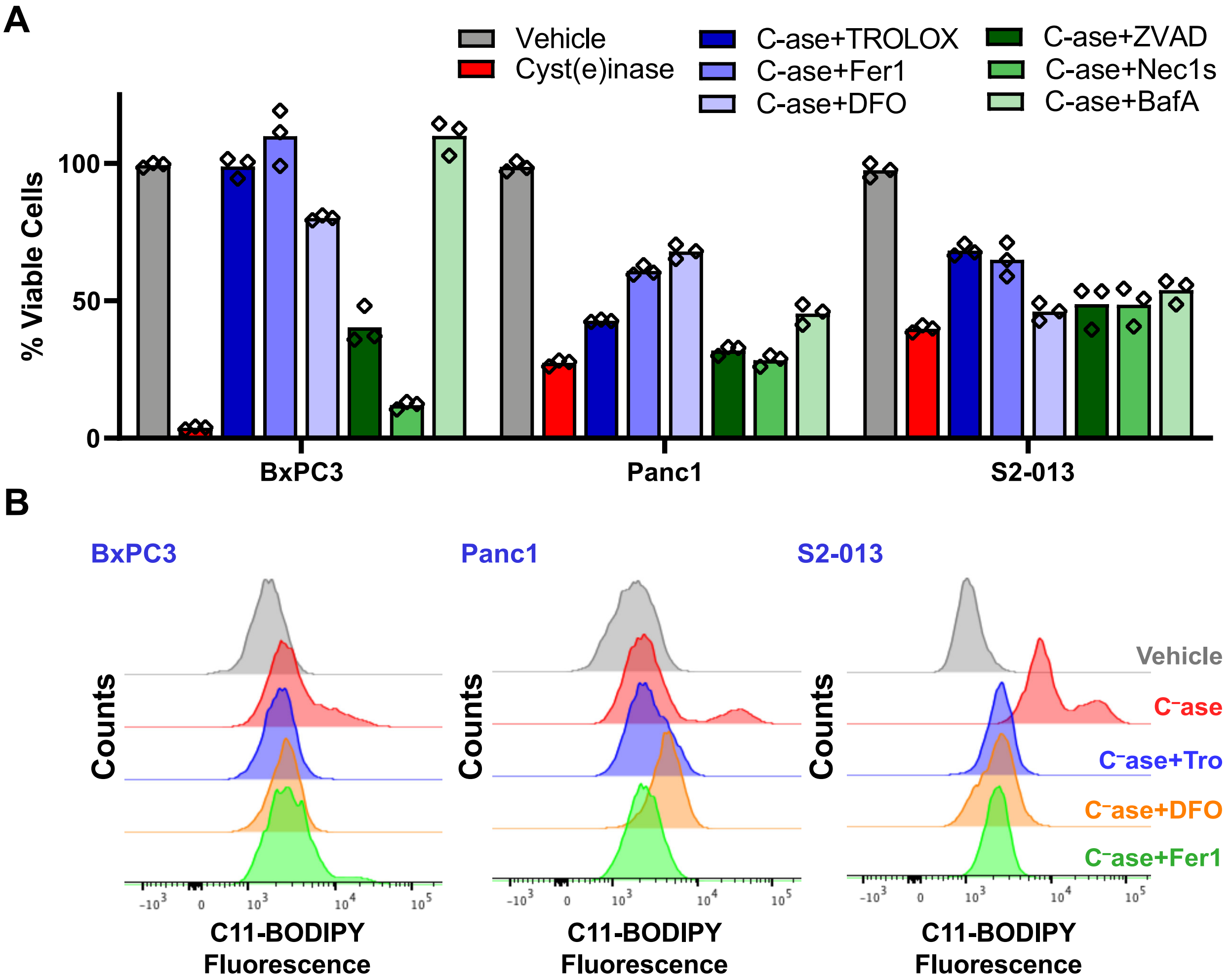


**Supplementary Figure 10. Cysteinase in vitro studies. (A)** Viability of PDA cells treated with cyst(e)inase, alone or in combination with Trolox, Fer1, DFO, ZVAD-FMK, Necrostatin-1S, or Bafilomycin A. **(B)** C11-BODIPY fluorescence for PDA cells treated with cyst(e)inase alone or in combination with Trolox, DFO, or Fer1.
