## Supplementary Figure 11 for "“Induction of pancreatic tumor-selective ferroptosis through modulation of cystine import”"

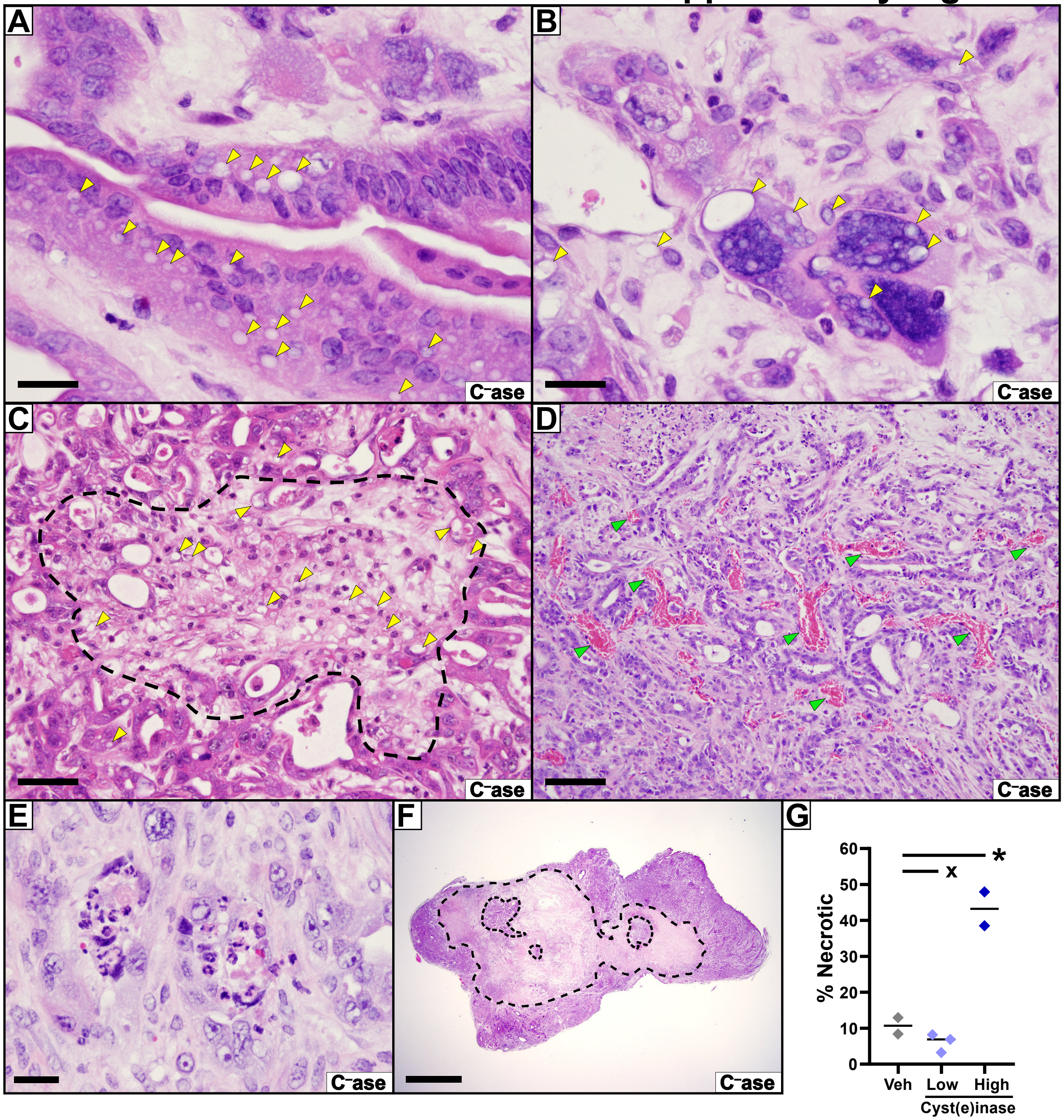


**Supplementary Figure 11. Histopathology of cyst(e)inase treated KPC pancreatic tumors. (A-F)** H&E stained microscopic images of pancreatic tumor tissues treated from KPC mice treated with cyst(e)inase. Yellow arrowheads show lipid droplets. (A,B) Images of well- (A) and poorly- (B) differentiated PDA treated with cyst(e)inase and exhibiting lipid droplet formation. Bars= 20µm. (C) Focal lesion (hashed line) exibiting large numbers of lipid droplets. Bar= 50µm. (D) In some cyst(e)inase tumors, decompressed blood vessels (green arrows) were noted. Bar= 100µm. (E) Scattered focal necrosis surrounded by viable tissue was noted in some tumors. (Bar= 20µm). (F) Regional necrosis (hashed line) of varying degrees was noted in all cyst(e)inase treated KPC tumors. (Bar= 2mm). **(G)** Quantification of necrotic regions in various treatment samples.
