## Supplementary Figure 12 for "“Induction of pancreatic tumor-selective ferroptosis through modulation of cystine import”"

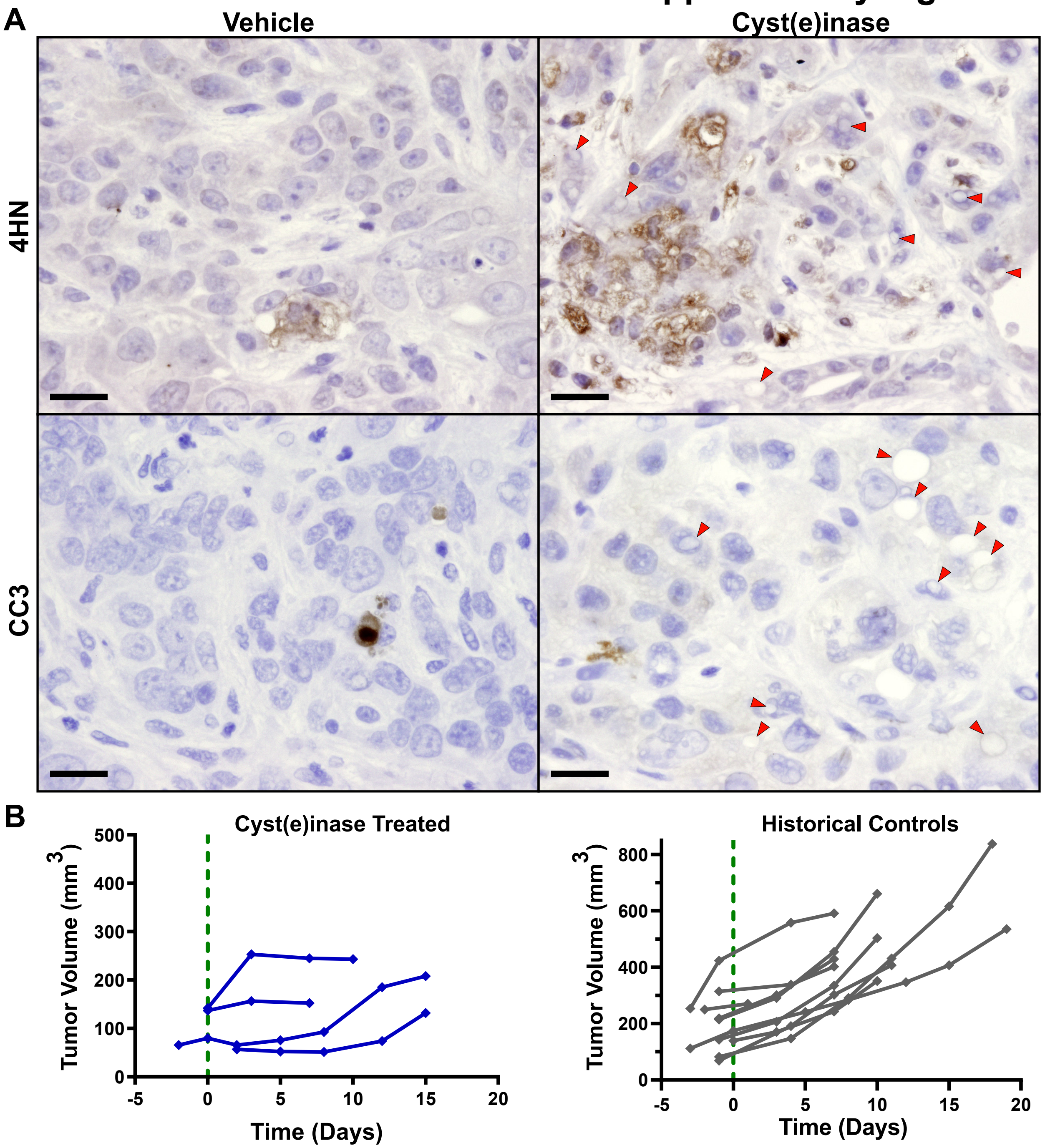


**Supplementary Figure 12. Response of KPC pancreatic tumors to cyst(e)inase treatment. (A)** Immunohistochemistry for 4-hydroxynonenol and cleaved caspase 3 from representative KPC pancreatic tumors treated with vehicle or cyst(e)inase in the 10-day short term response study. Red arrows indicate lipid droplets. **(B)** Tumor volumes from 3D high resolution ultrasound of KPC pancreatic tumors treated and imaged longitudinally. Four tumors from three cyst(e)inase-treated mice show evidence of altered growth kinetics in response to treatment. By contrast, tumors from historical control animals have never shown spontaneous tumor stabilizations or regressions.
