## Supplementary Video Legends for "“Induction of pancreatic tumor-selective ferroptosis through modulation of cystine import”"

Supplemental Video 1. PANC-1 cells die from cystine withdrawal.

Here, we present a 20 fps video of PANC-1 cells cultured in the absence of cystine. Each frame is captured per minute of real time (thus, each second of video is 20 minutes of real-time) In this video, cells are imaged from hours 6-12 of treatment and begin to exhibit evidence of catastrophic membrane damage beginning at about 8 hours and concluding by 12 hours.

Supplemental Video 2. PANC-1 cells die from system x_c_^-^ inhibition.

Here, we show time-lapse microscopy of PANC-1 cells treated with 1 µM IKE for the duration of the experiment. Once again, one frame was captured per minute of real time. The video begins after 6 hours of treatment and ends at approximately 12 hours. Similarly to cystine withdrawal, these cells die from membrane ballooning effects in a similar time span.

Supplemental Video 3. Staurosporine causes apoptosis in PANC-1 cells and is morphologically distinct from cysteine deprivation-induced cell death.

Time-lapse microscopy of PANC-1 cells treated with .2 µM staurosporine from 6-12 hours. Staurosporine is a known apoptosis inducer and here causes the nuclear fragmentation phenotype typical of this process. This, however, is very distinct from the cell death caused by cysteine deprivation, as seen in previous videos. Capture details are akin to previous videos.
